# One mucosal administration of a live attenuated recombinant COVID-19 vaccine protects nonhuman primates from SARS-CoV-2

**DOI:** 10.1101/2021.07.16.452733

**Authors:** Mariana F. Tioni, Robert Jordan, Angie Silva Pena, Aditya Garg, Danlu Wu, Shannon I. Phan, Xing Cheng, Jack Greenhouse, Tatyana Orekov, Daniel Valentin, Swagata Kar, Laurent Pessaint, Hanne Andersen, Christopher C. Stobart, Melissa H. Bloodworth, R. Stokes Peebles, Yang Liu, Xuping Xie, Pei-Yong Shi, Martin L. Moore, Roderick S. Tang

**Author notes:** Joint first authors (Mariana F. Tioni, Robert Jordan). Corresponding author: Mariana Tioni, Meissa Vaccines Inc, 1100 Island Drive, Ste 202, Redwood City, CA 94065.

## Abstract

Severe Acute Respiratory Syndrome Coronavirus 2 (SARS-CoV-2) is the causative agent of the COVID-19 global pandemic. SARS-CoV-2 is an enveloped RNA virus that relies on its trimeric surface glycoprotein, spike, for entry into host cells. Here we describe the COVID-19 vaccine candidate MV-014-212, a live attenuated, recombinant human respiratory syncytial virus (RSV) expressing a chimeric SARS-CoV-2 spike as the only viral envelope protein. MV-014-212 was attenuated and immunogenic in African green monkeys (AGMs). One mucosal administration of MV-014-212 in AGMs protected against SARS-CoV-2 challenge, reducing by more than 200- fold the peak shedding of SARS-CoV-2 in the nose. MV-014-212 elicited mucosal immunity in the nose and neutralizing antibodies in serum that exhibited cross-neutralization against two virus variants of concern. Intranasally delivered, live attenuated vaccines such as MV-014-212 entail low-cost manufacturing suitable for global deployment. MV-014-212 is currently in phase 1 clinical trials as a single-dose intranasal COVID-19 vaccine.

## Introduction

<u>CO</u>rona<u>VI</u>rus <u>D</u>isease 20<u>19</u> (COVID-19) is the latest virus pandemic to afflict humanity. The disease began its spread in the late months of 2019^1, 2^, and by March 11, 2020, 118,000 people across 114 countries were infected, at which time it was declared a pandemic by the World Health Organization (WHO)^3^. COVID-19 is a respiratory disease often leading to pneumonia, caused by the highly transmissible Severe Acute Respiratory Syndrome Coronavirus 2^4^. The overall mortality rate is approximately 2% and the disease is especially severe in the elderly and in patients with serious underlying medical conditions, such as heart or lung disease and diabetes. As of July 30, 2021, there were 193,553,009 confirmed cases of SARS-CoV-2 infection with a total of 4,200,412 deaths worldwide (WHO dashboard, https://covid19.who.int/).

SARS-CoV-2 is an enveloped RNA virus that relies on its surface glycoprotein, spike, for entry into host cells^5, 6^. The spike protein is a type I fusion protein that forms a trimer that protrudes on the viral membrane, giving the virus its characteristic crownlike appearance under electron microscopy^7, 8^. The angiotensin-converting enzyme 2 (ACE2) has been identified as a cellular receptor for SARS-CoV-2 spike^5, 9^. Disrupting the interaction of ACE2 and the receptor binding domain (RBD) of spike is at the core of vaccine design and therapeutics. Currently, three COVID-19 vaccines are approved for emergency use in the United States^10^. The three vaccines are based on the SARS-CoV-2 spike protein, and their high level of efficacy has validated spike as a protective antigen.

All the emergency use authorization (EUA) vaccines currently in use are delivered intramuscularly, and none is live attenuated. Live attenuated vaccines (LAVs) are often administered by the same route of entry as the pathogen they target, and LAVs replicate in the host, mimicking natural infection without causing disease. For respiratory viruses, intranasal LAVs generate mucosal immunity at the site of infection, blocking the pathogen at the earliest phases of infection thus helping control systemic spread^11^. In the case of influenza infection, LAV induces better mucosal immunoglobulin A (IgA) and cell-mediated immunity relative to other vaccine types, eliciting a longer-lasting and broader immune response that more closely resembles natural immunity^12^. Furthermore, comparison of intramuscular vs intranasal vaccines against SARS-CoV in mice showed that serum IgA was only induced following intranasal vaccination^13^, and only intranasal vaccination provided protection in both upper and lower respiratory tracts^14^. For SARS-CoV-2 in particular, the early antibody response is dominated by IgA and mucosal IgA is highly neutralizing^15^, underscoring the importance of developing an intranasal vaccine capable of eliciting mucosal immunity. According to the WHO vaccine tracker, there are currently 108 vaccines in clinical trials, only 4 of which are intranasal replicating vaccines^16^.

Here we describe the rational design, generation, and preclinical evaluation of a novel COVID-19 vaccine candidate. MV-014-212 is a live attenuated recombinant vaccine strain derived from the human respiratory syncytial virus (RSV) vaccine candidate OE4^17^. In MV-014-212, the RSV attachment (G) and fusion (F) surface glycoproteins were replaced with a chimeric SARS-CoV-2 spike harboring the cytoplasmic tail of the RSV F protein. MV-014-212 was attenuated and immunogenic in nonhuman primates, producing both systemic and mucosal immunity after one mucosal administration. A single mucosal administration of MV-014-212 in African green monkeys (AGMs) elicited neutralizing antibodies against the homologous virus and cross-neutralizing activity against the variants of concern, B.1.351 and B.1.1.7. Studies in mice indicated that MV-014-212 vaccination generated a Th1-biased cellular immune response. MV-014-212 is currently being evaluated in phase 1 clinical trials as a single-dose intranasal COVID-19 vaccine.

## Results

### Design and Generation of MV-014-212

MV-014-212 is a novel, live attenuated, recombinant vaccine against SARS-CoV-2, based on the backbone of the human respiratory syncytial virus (RSV) (Fig. 1). The G and F proteins of RSV were replaced by a chimeric protein consisting of the ectodomain and transmembrane (TM) domains of SARS-CoV-2 spike (USA-WA1/2020) and the cytoplasmic tail of RSV F (strain line 19). The rational design of this chimera was based on previous observations of the need for the homologous TM of spike in coronavirus infectious particle production^18–20^ and the reported importance of the cytoplasmic tail of RSV F in the production of RSV progeny^21^. The sequence of amino acids at the junction between spike and F proteins is shown in Fig. 1. Notably, the chimeric spike/RSV F protein retains functionality as MV-014-212 growth relies on it for attachment and fusion with the host cell. Moreover, since this is the only surface protein of MV-014-212, the possibility of negative interference due to preexisting immunity against RSV is eliminated. Various chimeric spike constructs that differed in the junction position were assessed for growth in Vero cells (Supplementary Fig. S1). In particular, a construct with the entire native SARS-CoV-2 spike was evaluated (MV-014-300, Fig. S1). While this construct could be rescued, it did not propagate productively in cell culture, demonstrating that the cytoplasmic tail of the F protein contributes to viral growth of the chimeric virus expressing spike. Of the constructs expressing different chimeric spike/RSV F fusion proteins, MV-014-212 was selected for further evaluation based on the ease of rescue and its ability to grow to acceptable titers for preclinical and clinical studies.

**Fig. 1:**
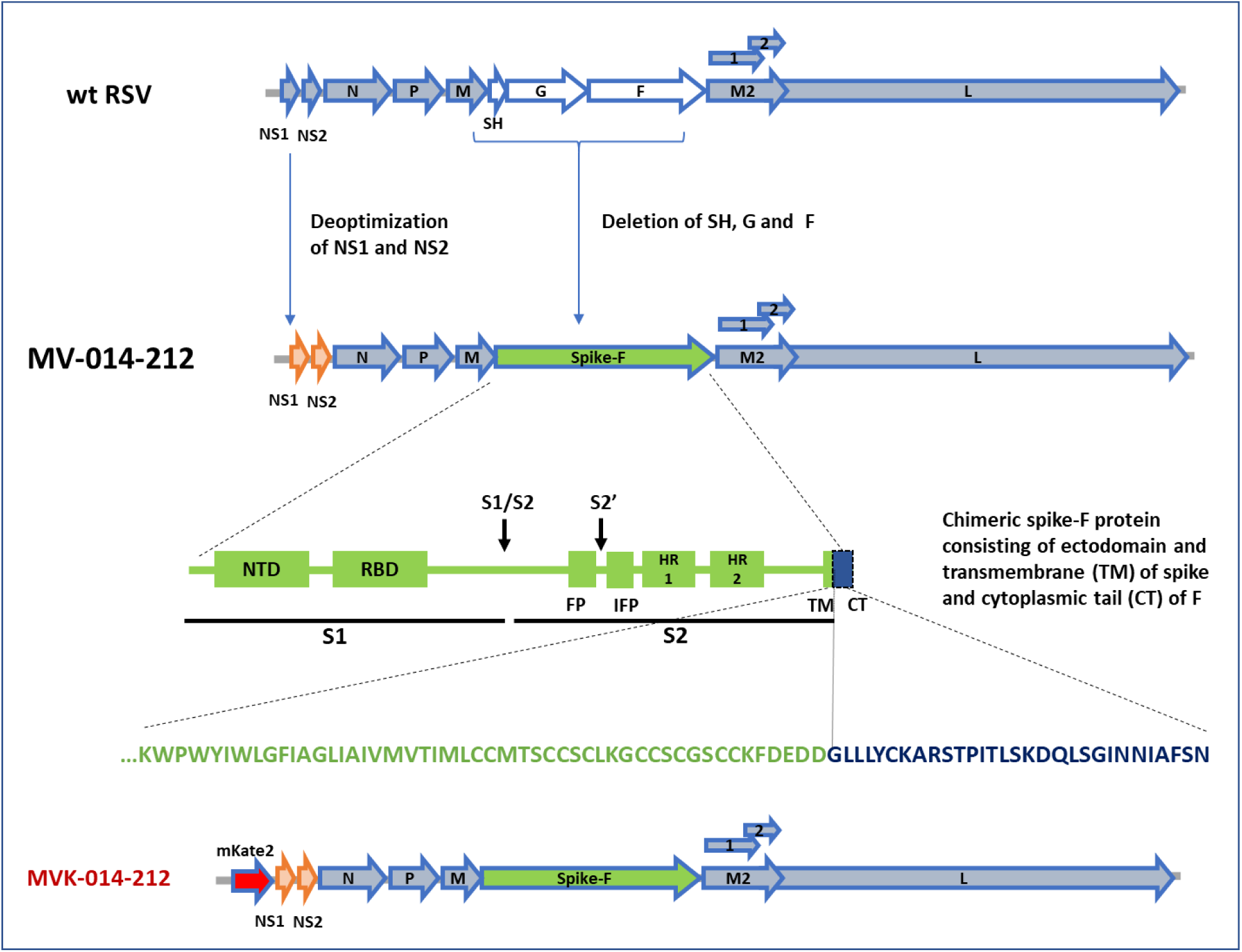
Design of MV-014-212. In MV-014-212, the *NS1* and *NS2* genes are deoptimized and the RSV *SH*, *G*, and *F* genes are deleted and replaced by a gene encoding a chimeric protein spike-F. The amino acid sequence at the junction is shown below the block graphic. The transmembrane domain of spike is represented in green, and the cytoplasmic tail of F is depicted in blue. The reporter virus MVK-014-212, encoding the fluorescent protein mKate2 in the first gene position, is schematically shown at the bottom or the panel. CT, cytoplasmic tail; FP, fusion peptide; IFP, internal fusion peptide; HR1 and 2, heptad repeats 1 and 2; NTD, N-terminal domain; RBD, receptor binding domain; S1, subunit S1; S2, subunit S2; S1/S2 and S2’, protease cleavage sites; TM, transmembrane domain.

The RSV backbone used to generate MV-014-212 was attenuated for replication in primary cells by codon deoptimization of the genes encoding the proteins NS1 and NS2 that suppress host innate immunity^22^. In addition, the short hydrophobic glycoprotein SH was deleted (Fig. 1) to further attenuate the virus in vivo and increase transcription of downstream genes^17^.

To facilitate the development of a microneutralization assay, a reporter virus derived from MV-014-212 was constructed by inserting the gene encoding the fluorescent mKate2 protein^23, 24^ upstream of the *NS1* gene (MV<u>K</u>-014-212, Fig. 1).

The recombinant virus constructs were electroporated into Vero cells and infectious virus was rescued and propagated for further characterization^24^. In MV-014-212, CPE is observed as the formation of polynucleated bodies or syncytia and eventual cell detachment (Fig. 2a). The electroporated cells were expanded until the CPE was extensive and the virus stock was harvested as a total cell lysate. The titers obtained for MV-014-212 and MVK-014-212 were comparable and within the range 1-5x10^5^ PFU/mL.

**Fig. 2:**
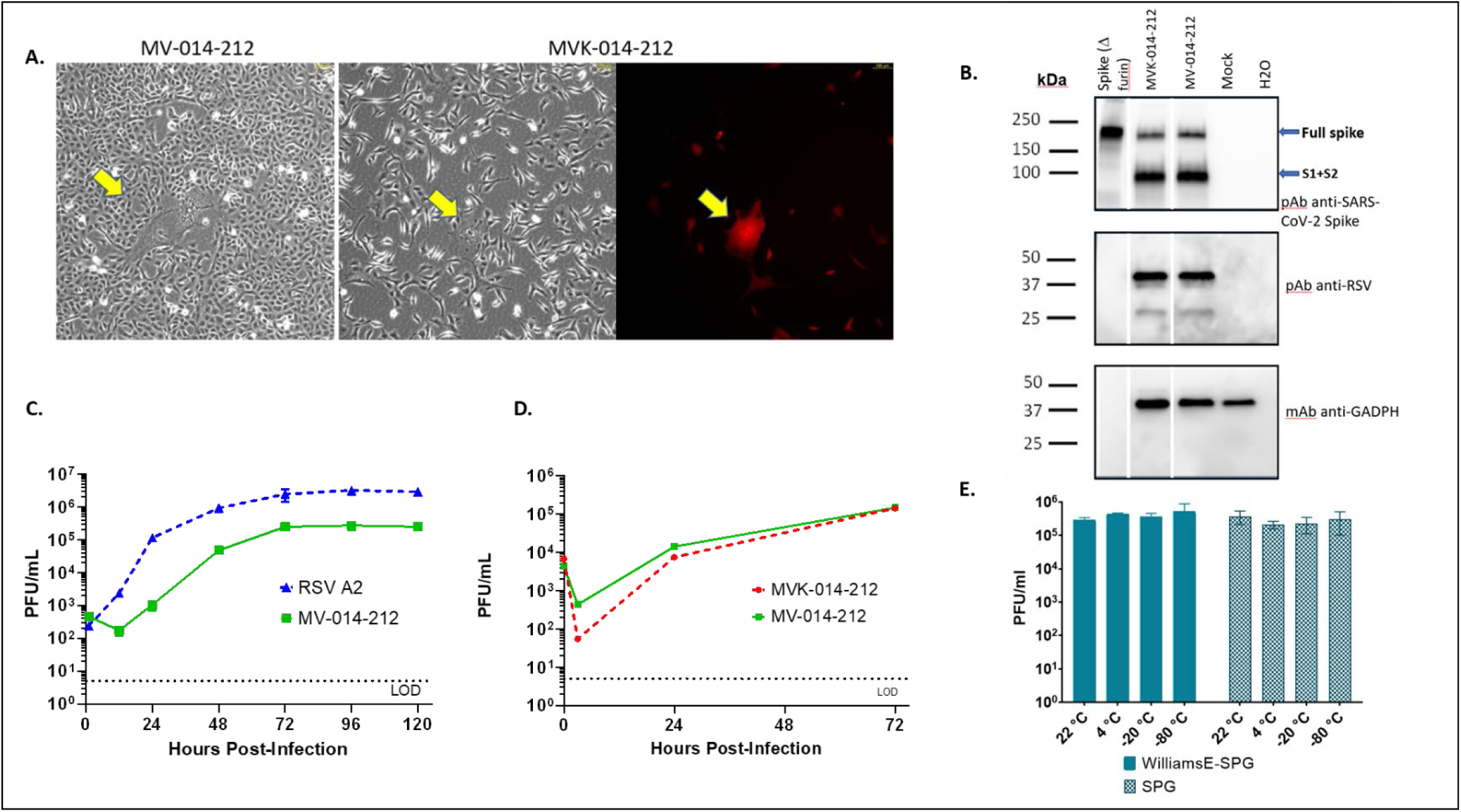
In vitro characterization of MV-014-212 a. Syncytia formed by MV-014-212 and MVK-014-212. Micrographs were taken at a total amplification of 100x under phase contrast or using tetramethylrhodamine (TRITC) filter. Yellow arrows point to syncytia. **b.** Western blot showing full-length purified SARS-CoV-2 spike protein lacking the furin cleavage site (LakePharma, lane 1), MVK-014-212 (lane 2), MV-014-212 (lane 3), mock-infected Vero cell lysate (lane 4), blank, (water, lane 5). The molecular weight marks correspond to the migration of the BIO-RAD Precision Plus Protein Dual Color Standards. The blue arrows indicate the expected migration of the full-length spike and the cleaved protein (S1+S2). **c** Multicycle replication kinetics of MV-014-212 compared to RSV A2 in serum-free Vero cells. Cells were infected at an MOI of 0.01 and incubated at 32 °C. Cells and supernatants were collected at 0, 12, 24, 48, 72, 96, and 120 hours post infection. Titers of the samples were determined by plaque assay in Vero cells. Data points represent the means of 2 replicate wells and error bars represent the standard deviation. **d** Multicycle replication kinetics of MV-014-212 compared with MVK-014-212 in serum-free Vero cells. Cells were infected at an MOI of 0.01 and incubated at 32 °C. Cells and supernatants were collected at 0, 3, 24, and 72 hours post infection. Titers of the samples were determined by plaque assay in Vero cells. Data points represent the means of 3 replicate wells and error bars represent the standard deviation. **e** Short-term thermal stability assay. Virus stocks of MV-014-212 prepared in Williams E medium supplemented with sucrose phosphate glutamate buffer (SPG) or prepared in SPG alone were incubated for 6 hours at –80 °C, 4 °C, –20 °C, and room temperature and the titer after incubation was determined by plaque assay. MOI, multiplicity of infection.

### In vitro characterization of MV-014-212

The SARS-CoV-2 spike protein contains a cleavage site between the S1 and S2 domains that is processed by furin-like proteases^25^ (Fig. 1). As for other coronaviruses, the S1 and S2 subunits of SARS-CoV-2 spike are believed to remain noncovalently bound in the prefusion conformation after cleavage^26, 27^. To determine if the chimeric spike protein encoded by MV-014-212 is expressed and proteolytically processed, virus stocks prepared from lysates of infected Vero cells were analyzed on western blots and probed with polyclonal antiserum against SARS-CoV-2 spike protein. Both MV-014-212 and MVK-014-212 viruses express the full length and cleaved forms of the chimeric spike protein (Fig. 2b), consistent with partial cleavage at the S1-S2 junction, with the expected apparent sizes^28, 29^ (Supplementary Fig. S2).

Multicycle growth kinetics of MV-014-212 was compared to wild-type (*wt)* recombinant RSV A2 in Vero cells (Fig. 2c). MV-014-212 exhibited delayed growth kinetics relative to RSV A2, showing an initial lag phase of approximately 12 hours. Both viruses reached their peak titers at 72 hpi and the titers remained constant until 120 hpi. The peak titer for MV-014-212 was approximately one order of magnitude lower than that of RSV A2. To determine if the insertion of the *mKate2* gene affected replication kinetics of MVK-014-212, Vero cells were infected with MV-014-212 or MVK-014-212. The growth kinetics of MVK-014-212 was similar to that of MV-014-212, reaching comparable peak titers by 72 hpi (Fig. 2d). These data are consistent with a report that insertion of *mKate2* in the first gene position did not significantly attenuate RSV A2- line 19F in vitro^24^.

To evaluate the short-term thermal stability of MV-014-212, aliquots of the viral stock were incubated at different temperatures for a period of 6 hours and the amount of infectious virus after the incubation was determined by plaque assay. Two stocks of MV-014-212 prepared in different excipients were compared in this study (Fig. 2e). The results demonstrate that MV-014-212 is stable for at least 6 hours in either excipient at –80 °C and room temperature. The genetic stability of MV-014-212 was examined by serial passaging in Vero cells. Subconfluent Vero cells were infected in triplicate with an aliquot of MV-014-212 and passaged for 10 consecutive passages (Supplementary Fig. S3). Viral RNA was isolated from passages 0 and 10 and amplified by reverse transcription polymerase chain reaction (RT-PCR). The sequence of the entire coding regions of the viral genome was determined by Sanger sequencing. The results showed that for all three lineages there were no variations detected at passage 10 relative to the starting stock (passage 0). Thus, the vaccine candidate was highly genetically stable in vitro.

### MV-014-212 replication is attenuated in African green monkeys and confers protection against *wt* SARS-CoV-2 challenge

African green monkeys (AGMs) are semi-permissive for replication of both *wt* SARS-CoV-2^30–33^ and RSV^34^, and therefore constitute an appropriate nonhuman primate model for studying the attenuation and protective immunogenicity of MV-014-212.

The AGM study design is depicted in Fig. 3a. On Day 0, AGMs were inoculated via the intranasal (IN) and intratracheal (IT) routes with 1.0 mL of 3×10^5^ PFU/mL MV-014-212 or *wt* RSV A2 at each site for a total dose of 6×10^5^ PFU per animal. Due to the semi-permissive nature of the AGM model, IT inoculation was necessary to promote replication of the vaccine and the SARS-CoV-2 challenge virus in the lungs. Animals in the mock group were similarly mock-inoculated with phosphate-buffered saline (PBS). Nasal swabs (NS) and bronchoalveolar lavage (BAL) samples were collected through Day 12 after immunization. Viral shedding in NS and BAL samples was determined by plaque assay using fresh samples that were not frozen at the study site. The results showed that the level of infectious virus in animals inoculated with MV-014-212 and duration of shedding in nasal secretions were lower than in animals inoculated with RSV A2 (Fig. 3b and Supplementary Fig. S4). The mean peak titer for RSV A2 was approximately 20-fold higher than that observed for animals inoculated with MV-014-212. These results show that MV-014-212 is attenuated in the upper respiratory tract of AGMs, compared to RSV A2.

**Fig. 3:**
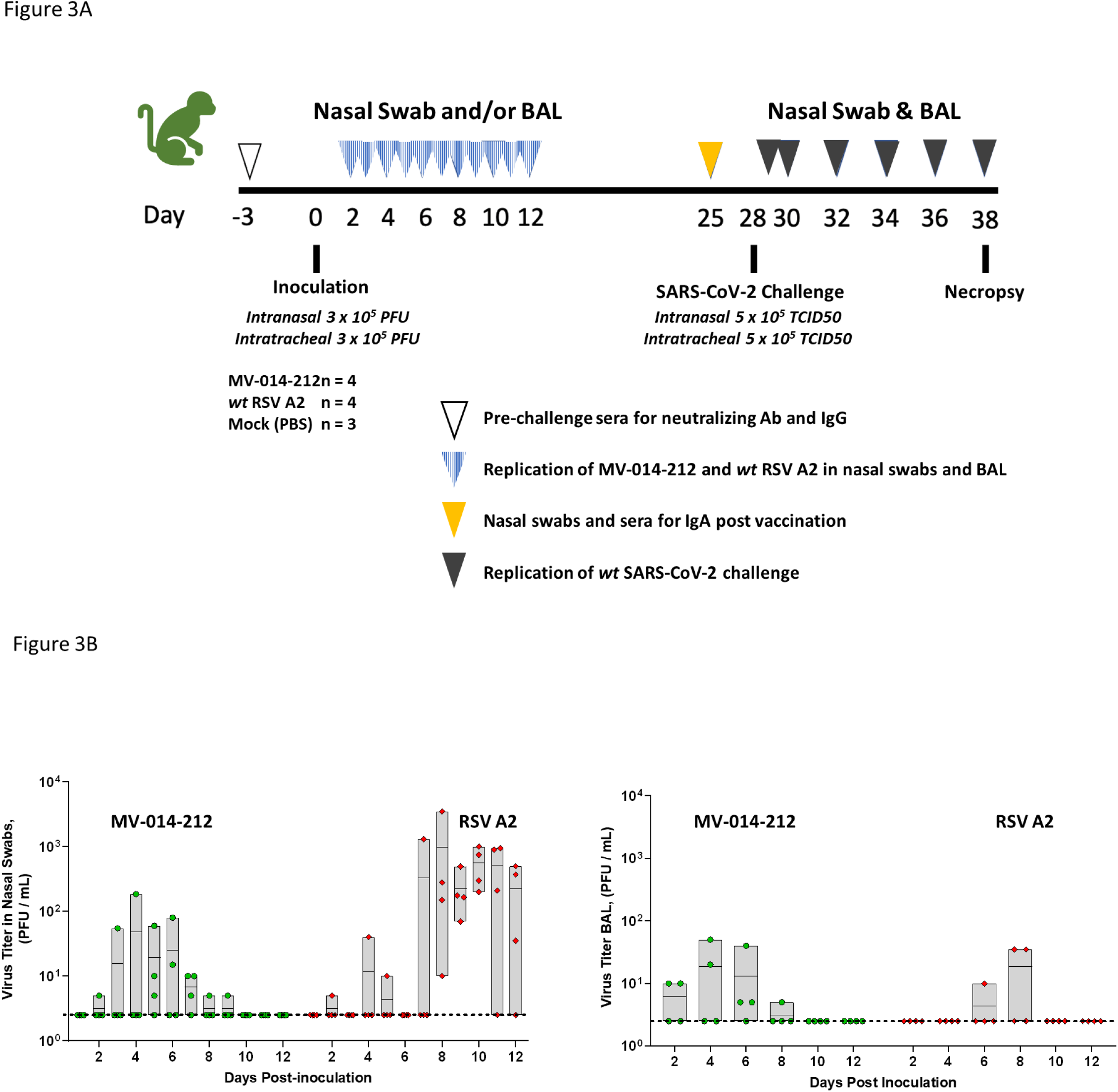
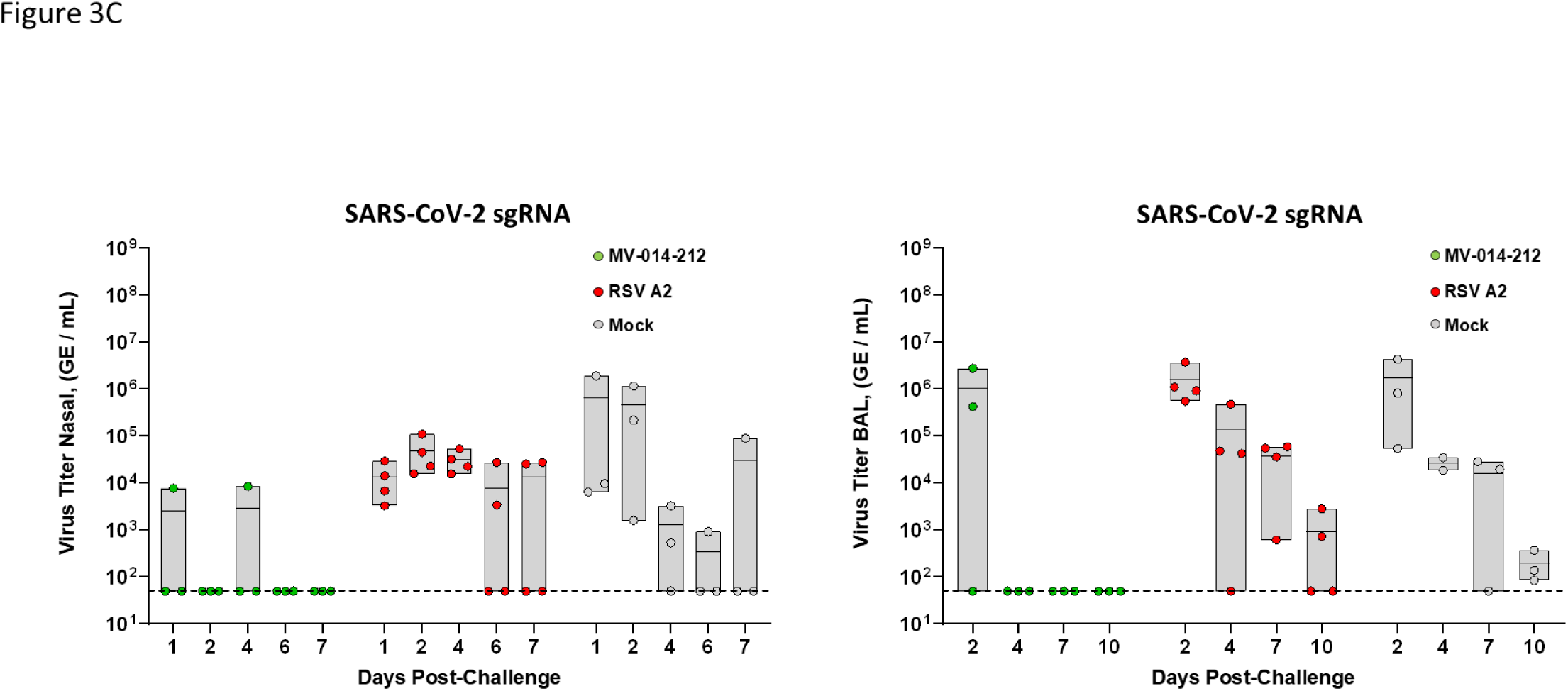
MV-014-212 attenuation in AGMs and protection against SARS-CoV-2 challenge a. AGM Study design. Nasal swabs (NS) were obtained on Days 1 through 12. Bronchoalveolar lavage (BAL) were collected on Days 2, 4, 6, 8, 10, and 12. Viral shedding in NS and BAL samples was determined by plaque assay using fresh samples. On Day 28 post inoculation AGMs were challenged with *wt* SARS-CoV-2. NS were obtained every day from Day 29 through 38. BAL was obtained on alternate days starting on Day 30 to Day 38. RT-qPCR was used for detecting SARS-CoV-2 shedding in NS and BAL samples. **b** Attenuation of MV-014-212 in the upper and lower respiratory tract of AGM. Viral titer in nasal swabs (A) or BAL (B) from AGMs following inoculation with MV-014-212 or *wt* RSV A2 were measured by plaque assay on Vero cells. On Days 1 through 12 post inoculation NS were collected in Williams E supplemented with SPG. Viral titer in BAL was measured on Days 2, 4, 6, 8, 10, and 12 post inoculation. The horizontal line in the box is the mean value of the data points for each time point. The dotted line represents the LOD (50 PFU/mL). For shedding kinetics of individual animals see Supplementary Fig. S5. **c** Protection of MV-014-212–vaccinated AGMs against *wt* SARS-CoV-2 challenge (sgRNA). *Wt* SARS-CoV-2 sgRNA in NS samples from AGMs inoculated with MV-014-212, *wt* RSV A2 or PBS (mock) following challenge. At Day 28, animals were challenged with 1.0 x 10^6^ TCID_50_ of *wt* SARS-CoV-2 by intranasal and intratracheal inoculation. NS collected on Days 1, 2, 4, and 6 post challenge with *wt* SARS-CoV-2 are shown. The level of SARS-CoV-2 sgRNA was determined by RT-qPCR. The dashed line represents the LOD of 50 genome equivalents (GE)/mL. For shedding kinetics of individual animals see Supplementary Fig. S6. AGM, African green monkey; BAL, bronchoalveolar lavage; LOD, limit of detection; PBS, phosphate-buffered saline; RT-qPCR, quantitative reverse transcription polymerase chain reaction; SPG, sucrose phosphate glutamate; TCID_50_, median tissue culture infectious dose; *wt*, wild type.

Low to undetectable virus titers were also observed in the lower respiratory tract of animals inoculated with MV-014-212 or RSV A2 over the course of 12 days. Both viruses replicated at low levels, but peak levels occurred earlier for MV-014-212. In this study, RSV A2 showed 100- to 1000-fold lower peak titers in the lower respiratory tract of AGMs compared with *wt* RSV A2 titers reported in literature^35–38^, confounding the ability to assess attenuation of MV-014-212 in the lungs. Subsequently, lower titers were also observed in the lungs of cotton rats for the recombinant A2 used in this study (rA2 from Meissa Vaccines Inc., Supplementary Fig. S5), relative to biologically derived RSV strains, suggesting that the RSV A2 used in this AGM study was attenuated in lungs.

Nasal and BAL samples from Day 6 post vaccination were used to extract RNA for sequence analyses of the spike gene of MV-014-212. Using Sanger sequencing, no variations in the spike gene were detected compared to the reference sequence for MV-014-212.

On Day 28 AGMs were challenged with 10^6^ median tissue culture infectious dose (TCID_50_) of *wt* SARS-CoV-2. NS and BAL samples were collected for 10 days after challenge. Shedding of *wt* SARS-CoV-2 was measured by quantitative RT-PCR (RT-qPCR) of the *E* gene subgenomic SARS-CoV-2 RNA (sgRNA) (Fig. 3c and Supplementary Fig. S6).

MV-014-212–vaccinated monkeys had low or undetectable levels of *wt* SARS-CoV-2 sgRNA in NS samples, in contrast to animals inoculated with *wt* RSV A2 or PBS (mock), which had higher levels of SARS-CoV-2 sgRNA. While the level of SARS-CoV-2 sgRNA was undetectable in animals vaccinated with MV-014-212 at most time points, one animal had detectable SARS-CoV-2 sgRNA at Day 2 and a different animal had a similar titer at Day 4 post challenge (Supplementary Fig. S6). Mean peak titers of SARS-CoV-2 in NS of animals in the control RSV and PBS groups were 20- and 250-fold higher than for animals vaccinated with MV-014-212, respectively. In both RSV- and mock-infected animals, shedding of *wt* SARS-CoV-2 sgRNA decreased steadily in nasal secretions from Days 4 to 10, and by Day 10 all animals in both groups had undetectable SARS-CoV-2 sgRNA.

Vaccination with MV-014-212 resulted in increased clearance of SARS-CoV-2 in lungs compared to RSV A2 or mock inoculation with PBS (Fig. 3c and Supplementary Fig. S6). The peak titer of SARS-CoV-2 in BAL samples occurred at Day 2 and was similar in all three treatment groups. Lung titers were undetectable in MV-014-212–vaccinated animals on Days 4 through 10 whereas SARS-CoV-2 was readily measured in animals inoculated with RSV A2 or PBS. Shedding of infectious SARS-CoV-2 in BAL and NS after challenge was quantified by a TCID_50_ assay (Supplementary Fig. S7). The results show that AGMs vaccinated with MV-014-212 had on average between 100- and 1000-fold less infectious SARS-CoV-2 in BAL, relative to RSV A2 and PBS controls, at peak shedding days. The kinetics of clearance of the MV-014-212 group was also faster, in accordance with the results obtained by sgRNA qPCR assay. In NS, the average peak shedding of the MV-014-212 group was more than 1000-fold lower than that of the RSV A2 group (Fig. S7). In the NS of the mock-vaccinated group, however, one of the animals did not have detectable shedding of infectious virus at any of the time points (Fig. S7). As a result, the overall average peak shedding was only 30-fold higher in the mock group than that seen in the MV-014-212 vaccinated group.

Taken together, these data show that a single mucosal administration of MV-014-212 protected AGMs from *wt* SARS-CoV-2 challenge.

### Antibody responses to MV-014-212 vaccination in AGMs

SARS-CoV-2 spike-specific serum immunoglobulin G (IgG) and nasal IgA were measured by enzyme-linked immunosorbent assay (ELISA) in sera and NS, respectively, from AGMs immunized with MV-014-212, RSV A2, or PBS on Day 25 post immunization. All animals were seronegative for RSV A2 and SARS-CoV-2 at the start of the study. AGMs inoculated with MV-014-212 produced higher levels of SARS-CoV-2 spike-specific IgG in serum compared to AGMs inoculated with RSV A2 or PBS, which had levels of spike-specific IgG that were close to the limit of detection (Fig. 4a).

**Fig. 4:**
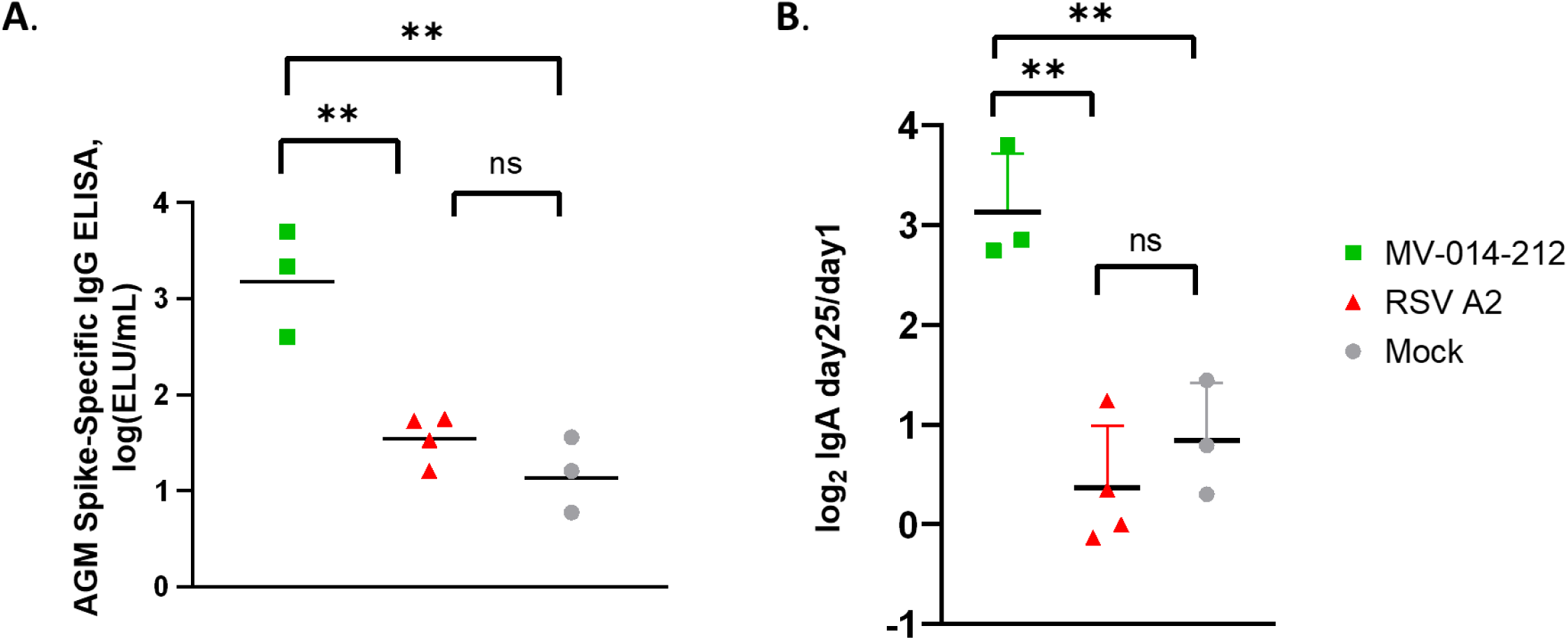
Spike-specific antibody responses in MV-014-212 inoculated AGMs. a. Spike-specific serum IgG antibodies specific to SARS-CoV-2 spike protein were measured by ELISA using serum collected on Day 25 from AGM inoculated with MV-014-212, *wt* RSV A2, or PBS (mock). The titer is expressed as ELISA units (ELU)/mL that were calculated by comparison against a standard curve generated from pooled human convalescent serum. **b** Spike-specific mucosal IgA. IgA antibodies specific to SARS-CoV-2 spike protein were measured by ELISA using NS collected on Day 25 post inoculation. The Log2 of the ratio of the values obtained at Day 25 over Day 1 are shown. The calculated ELU/mL concentration was obtained from a standard curve generated from total purified human IgA using a capture ELISA. Statistical analysis was one-way ANOVA. \*\**p* < 0.01. AGM, African green monkey; ANOVA, analysis of variance; ELISA, enzyme-linked immunosorbent assay; Ig, immunoglobulin; NS, nasal swabs; PBS, phosphate-buffered saline; *wt*, wild type.

Spike-specific IgA was also detected in the nasal swabs of monkeys inoculated with MV-014-212. There was more than an 8-fold increase in nasal spike-specific IgA in the MV-014-212– vaccinated animals 25 days after vaccination (Fig. 4b). In contrast, RSV A2 or mock-vaccinated animals did not show a significant change in spike-specific IgA.

These results showed that mucosal inoculation of MV-014-212 induced both nasal and systemic antibody responses to the functional SARS-CoV-2 spike.

To determine if vaccination with MV-014-212 elicited neutralizing antibodies in AGMs, three different approaches were used: (a) microneutralization assays with a fluorescent reporter homologous virus (MVK-014-212), (b) vesicular stomatitis virus (VSV) pseudovirus neutralization assays, and (c) plaque reduction assay with SARS-CoV-2 viruses. Fig. 5a shows a representative microneutralization assay where the fluorescent reporter MVK-014-212 was used to infect Vero cells in a 96-well plate. This assay was used to quantify the 50% neutralizing titers (NT_50_) of AGMs vaccinated with MV-014-212 or the control groups inoculated with RSV or PBS (Fig. 5b). The AGMs vaccinated with MV-014-212 showed an average NT_50_ of 99, whereas the RSV A2 and mock controls had an NT_50_ of 16 and 14, respectively. The same sera were used in a pseudovirus neutralization assay based on spike-pseudotyped vesicular stomatitis virus (VSV) particles carrying the luciferase reporter gene (Supplementary Fig. S8). The results of the VSV pseudovirion assay showed a similar trend but were not statistically significant, and there were higher background levels in the assay (Supplementary Fig. S8). The convalescent control is a pool of 3 human convalescent serum samples (Nexelis). Two animals in the MV-014-212 group with the higher serum nAb titers (Fig. 5b) were chosen to compare with this convalescent pool control. As with the pseudovirus assay (Supplementary Fig. S8), the microneutralization assay showed that the NT_50_ of the AGM sera were approximately 4-fold lower than the human convalescent pool (Fig. 5c). Lastly, the serum of one AGM vaccinated with MV-014-212 (the animal showing the highest titers in Fig. 5C), was used in a conventional 50% plaque-reduction neutralization assay with SARS-CoV-2 in BSL-3 containment^39, 40^. In this assay, the neutralizing titers (PRNT_50_) against SARS-CoV-2 virus or the two variants of concern, B.1.351 and B.1.17, were determined (Fig. 5d). The results showed that vaccination with MV-014-212 elicits neutralizing activity against the *wt* strain and both variants of concern (the pre-immune serum titers were below the limit of detection for all the virus strains tested, data not shown). The neutralization titer of B.1.351 was equivalent to that of the *wt* strain and B.1.1.7 showed slightly higher neutralizing titers. Taken together, the data suggested that MV-014-212 elicited modest serum neutralizing antibody titers in AGMs, and there was cross-neutralization against the variants B.1.351 and B.1.1.7.

**Fig 5:**
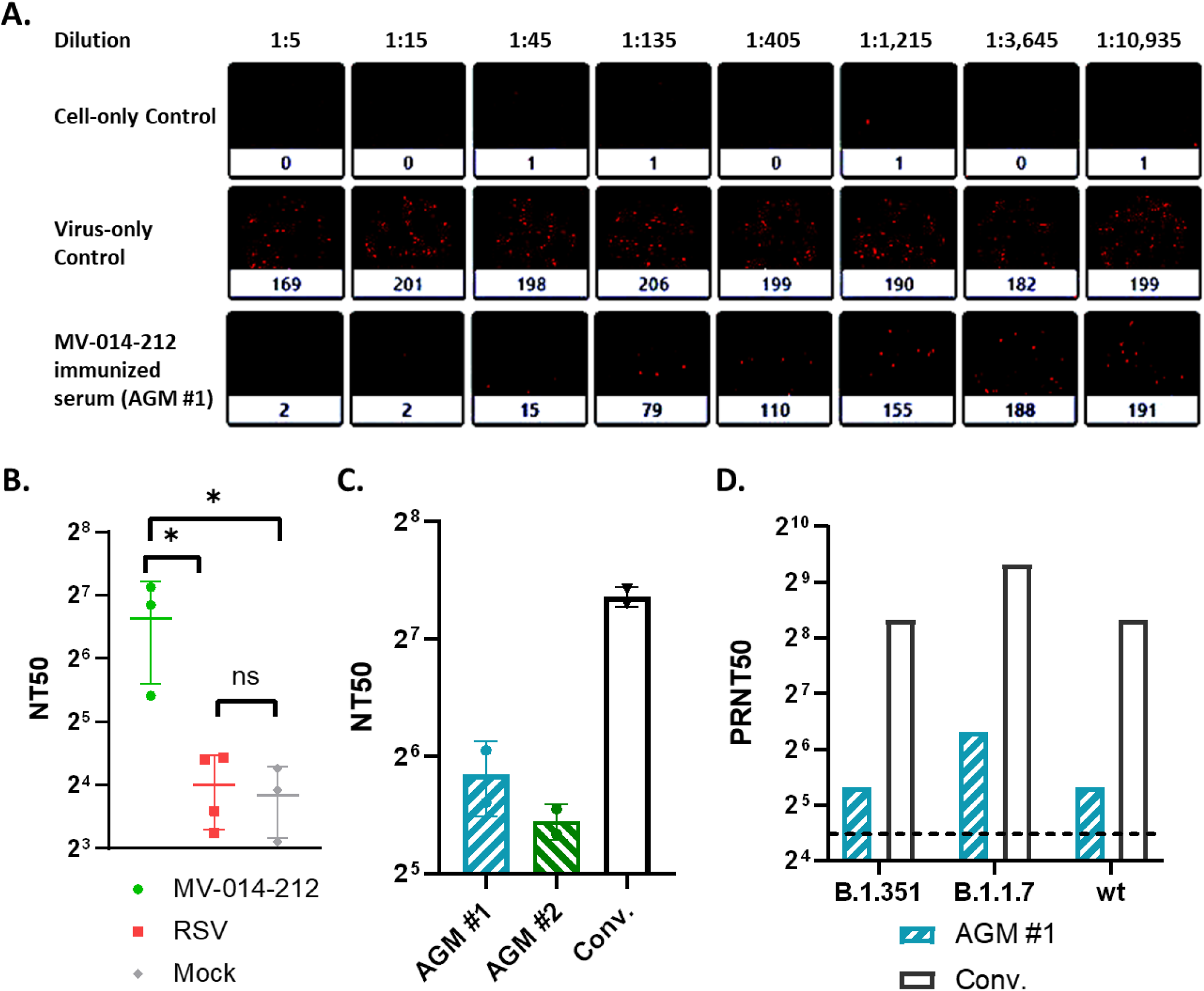
Neutralization assays with AGM sera. a. Representative microneutralization assay performed with the fluorescent reporter virus MVK-014-212. The top 2 rows are the cell-only and virus-only controls. The third row corresponds to the serum of an AGM immunized with MV-014-212 (AGM #1). The numbers on top represent the serum dilutions used and the number below each well is the count of viral plaques (that appear as red foci in the wells). The image of each well was captured with a Celigo Image Cytometer (Nexcelom) and the Celigo proprietary software was used to count the plaques. **b** Microneutralization assay using MVK-014-212. Neutralizing titers (NT_50_) in serum of AGMs immunized either with MV-014-212 or the negative controls, RSV or mock, obtained from a microneutralization assay (performed as in **a**). The plaque count from A was used to calculate the NT_50_ as described in Methods. **c** Microneutralization assay using MVK-014-212 comparing the NT_50_ of 2 AGM immunized with MV-014-212 with a convalescent control at 1859 IU/mL. The 2 animals used in this experiment correspond to the animals with highest titers in experiment **b**. **d** Plaque reduction neutralization assay. Serum from AGM #1 immunized with MV-014-212 was used in a 50% plaque-reduction neutralization test performed in a BSL-3 facility^39, 40^. The neutralization titers of AGM #1 against *wt* SARS-CoV-2 or 2 variants of concern (B.1.351 and B.1.1.7) are shown. The convalescent serum (“Conv.”) used in this assay is the same as in **c**. LOD is 20 and is indicated as a dashed line. In all cases, the data shown is the average of 2 technical replicates. Statistical analysis was one-way ANOVA. \**p* < 0.05. AGM, African green monkey; ANOVA, analysis of variance; LOD, limit of detection; RSV, respiratory syncytial virus; *wt*, wild type.

### MV-014-212 elicits a Th1-biased cellular immune response in hACE2-mice

Mouse models of vaccine-associated enhanced respiratory disease (VAERD) suggest that an imbalance in type 1 (Th1) and type 2 (Th2) T helper cell immunity with a skewing towards Th2 response contributes to enhanced lung pathology following challenge^41^. To assess the balance of Th1 and Th2 immunity generated after vaccination with MV-014-212, transgenic mice expressing human ACE2 receptor were inoculated with a single dose of MV-014-212 or PBS by the intranasal route. A control group received an intramuscular prime and boost vaccination with SARS-CoV-2 spike protein formulated in alum, which has been shown to skew immunity towards a Th2 response^42^. On Day 28, serum was collected to measure total spike-specific IgG, IgG2a, and IgG1 by ELISA. In addition, spleens were collected and the number of splenocytes expressing interferon-γ (IFNγ) or interleukin (IL)-5 were measured by ELISpot assay. The ratio of IgG2a/IgG1 and the ratio of cells producing IFNγ/IL-5 are indicators of Th1-biased cellular immune response^42, 43^.

The results showed that MV-014-212 induced spike-reactive splenocytes as measured by ELISpot assay (Fig. 6a). Importantly, MV-014-212 induced higher numbers of splenocytes expressing IFNγ relative to IL-5 when cell suspensions were stimulated with a spike peptide pool, suggesting that vaccination with MV-014-212 produced a Th1-biased immune response. The ratio of IFNγ–producing cells to IL-5–producing cells in the MV-014-212 group was more than one order of magnitude higher than in the group vaccinated with alum-adjuvanted spike protein (Fig. 6b). Consistent with the ELISpot data, the ratios of IgG2a/IgG1 detected in serum were higher in the animals vaccinated with MV-014-212 than the control group vaccinated with alum-adjuvanted spike (Fig. 6c and 6d). These data suggest that intranasal vaccination with live attenuated, recombinant MV-014-212 induced a Th1-biased antiviral immune response.

**Fig. 6:**
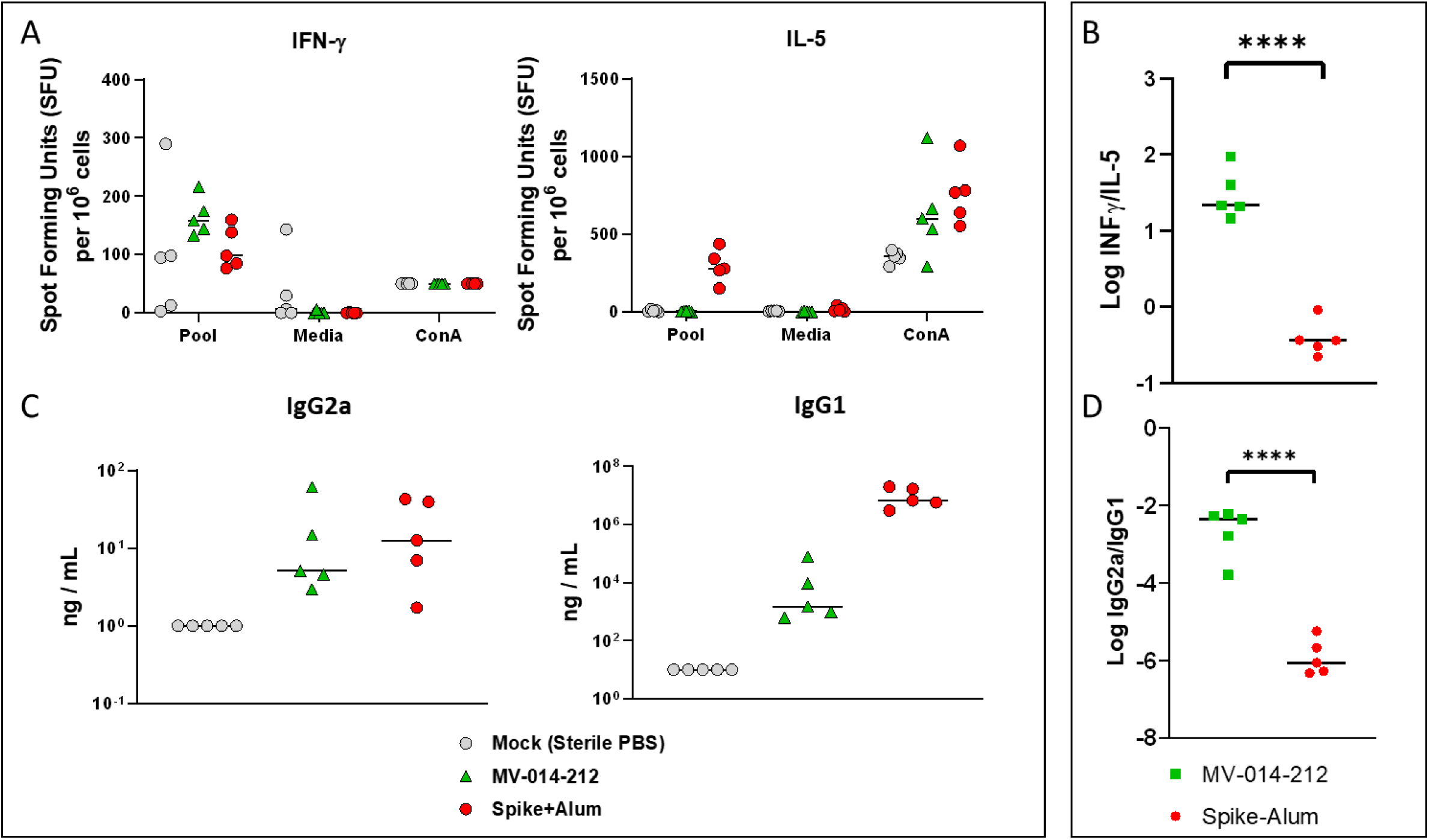
MV-014-212 elicited a Th1-biased immune response in ACE2 mice. a. ELISpot: IFNγ (left) or IL-5 (right)–producing cells in ACE2 mice. Splenocytes isolated from hACE-2–expressing mice (*n* = 5) were collected on Day 28 post inoculation (with MV-014-212 or PBS) and stimulated with a peptide pool that spanned the SARS-CoV-2 spike protein (pool), media, or the mitogen concanavalin A (Con A). IL-5 or IFNγ–expressing T cells were quantified by ELISpot assay. Control mice were vaccinated with purified SARS-CoV-2 spike protein adjuvanted with alum by intramuscular injection at Day –20 and Day 0. **b** Log of the ratio of IFNγ to IL-5–expressing cells (as shown in **a**.) **c** IgG1 and IgG2a ELISA. Levels of IgG2a (left panel) and IgG1 (right panel) corresponding to Day 28 serum from hACE-2 mice vaccinated intranasally with PBS or MV-014-212 or intramuscularly with spike-alum, as determined by ELISA. The concentration of each immunoglobulin isotype was determined from standard curves generated with purified SARS-CoV-2 spike specific monoclonal IgG2a or IgG1 antibodies. **d** Log of the ratio of IgG2a/IgG1 (as shown in **c**). Statistical analysis is an unpaired *t*-test. \*\*\*\**p* < 0.0001. ACE, angiotensin-converting enzyme; ELISA, enzyme-linked immunosorbent assay; hACE2, human ACE2; IFN, interferon; Ig, immunoglobulin; IL, interleukin; PBS, phosphate-buffered saline; Th, helper T cell.

## Discussion

MV-014-212 is a recombinant live attenuated COVID-19 vaccine designed to be administered intranasally to stimulate mucosal as well as systemic immunity against SARS-CoV-2. MV-014-212 was engineered to express a functional SARS-CoV-2 spike protein in place of the RSV membrane surface proteins F, G, and SH in an attenuated RSV strain expressing codon deoptimized *NS1* and *NS2* genes. Indeed, replication of MV-014-212 was attenuated in the respiratory tract of AGM following mucosal administrations in the nose and trachea and it elicited SARS-CoV-2 spike-specific mucosal IgA and serum IgG. Furthermore, vaccination with MV-014-212 induced a modest level of serum neutralizing antibodies and, more importantly, a single administration of the Meissa COVID-19 vaccine provided robust protection against SARS-CoV-2 challenge in AGM.

Based on the levels of viral sgRNA detected after SARS-CoV-2 challenge, the efficacy of MV-014-212 after one dose in nonhuman primates (NHPs) is comparable with that reported for the current EUA mRNA vaccines in NHPs. The efficacy and kinetics of clearance in MV-014-212– vaccinated AGMs are comparable to those observed after 2 doses of BNT126b2 or mRNA1273^44, 45^ in nasal swabs and BAL of rhesus macaques. hAdOx1 nCoV-19, on the other hand, confers comparable protection in the BAL but is less protective the in nose of rhesus monkeys^46^, compared with MV-014-212. The mRNA and other nonreplicating COVID vaccines were evaluated in rhesus monkeys, which are less permissive for RSV replication than AGMs^34^. AGMs were selected as the model for evaluating the attenuation and immunogenicity of MV-014-212 because they support both RSV^34^ and SARS-CoV-2^30–33^ replication. The level of SARS-CoV-2 detected in the mock-treated AGMs in this study was as high or higher than the levels reported for the mock groups in other vaccine studies^44–46^, allowing comparison of the results between different monkey species.

The use of a semi-permissive animal model for the evaluation of the LAV candidate MV-014-212 constitutes a limitation of this study, since LAVs rely on replication to elicit a robust immune response. The semi-permissiveness of AGMs to RSV and SARS-CoV-2 would result in lower replication of MV-014-212 and a more modest serum neutralizing antibody response. In a more permissive host, like humans, MV-014-212 is expected to replicate to higher titers, potentially resulting in higher immunogenicity than that observed in AGMs.

Remarkably, immunization of AGMs with MV-014-212 resulted in both mucosal and systemic antibody responses. There was approximately 100-fold more spike-specific total serum IgG in MV-014-212–vaccinated AGMs compared to AGMs receiving *wt* RSV A2 or PBS inoculations. Spike-specific IgA was also detected in nasal swabs of MV-014-212–immunized animals. There was approximately an 8-fold increase in IgA concentration 25 days following vaccination with MV-014-212. In an experimental human challenge study, low RSV F-specific mucosal IgA was a better predictor for susceptibility to RSV challenge in seropositive adults than serum IgG and neutralizing antibody levels^47^. Indeed, spike RBD-specific dimeric serum IgA was shown to be more potent at neutralizing SARS-CoV-2 than monomeric IgG^48^. By inference, secretory IgA, which exists at mucosal surfaces as dimeric IgA, may act as a potent inhibitor of SARS-CoV-2 at the site of infection. Interestingly, IgA antibodies dominate early humoral responses in human SARS-CoV-2 infections and IgA plasmablasts with mucosal homing potential peaked during the third week of disease onset^15^.

A SARS-CoV-2 neutralizing antibody response was detected in the serum of vaccinated AGMs in a microneutralization assay using an mKate2-expressing MV-014-212 virus (MVK-014-212). This response was also observed in a luciferase-based pseudovirus assay and in a conventional plaque reduction assay with SARS-CoV-2. The plaque reduction assay also detected a neutralizing response against 2 variants of concern, B.1.351 (Beta) and B.1.1.7 (Alpha), with comparable titers to the ones detected against the USA-WA2020 strain. Direct comparison of the AGM serum nAb titers to serum nAb titers published for nonreplicating vaccines in rhesus monkeys or to titers in human convalescent sera is confounded by the semi-permissivity of AGM for MV-014-212 replication. The immunogenicity of MV-014-212 is being studied in an ongoing phase 1 clinical trial. Peclinical data for two other intranasal vaccines were recently published^49, 50^.

The IM-administered EUA vaccines currently in use offer very satisfactory levels of protection, but their ability to prevent asymptomatic transmission is still unclear (reviewed by Tiboni et al^51^). A recent report from the Centers for Disease Control and Prevention (CDC)^52^ shows that in breakthrough infections, the viral load in the nose of vaccinated individuals is as high as that of unvaccinated patients. This finding and the rapid spread of the highly transmissible Delta variant underscore the urgent need for a vaccine that blocks infection in the nose and reduces transmission. Vaccines delivered intranasally stand a higher chance of eliciting local mucosal immunity, preventing not only systemic infection but also local replication and shedding. Indeed, IN-delivered MV-014-212 resulted in the production of spike-specific IgA in the noses of AGMs. According to the July 30, 2021 report^16^, “The landscape of candidate vaccines in clinical development,” prepared by WHO, there are 8 intranasal COVID-19 vaccines in clinical trials, only 2 of which are live attenuated viruses, MV-014-212 and COVI-VAC (Codagenix)^53^. Unlike COVI-VAC, MV-014-212 is a nonsegmented negative strand RNA virus not prone to recombine in nature. RNA recombination is extremely rare for nonsegmented negative strand RNA viruses outside of experimental co-infections in laboratory settings^54–56^. In addition to the lack of natural recombination, the results presented in this study showed that MV-014-212 was genetically stable. Neither accumulation of mutations nor loss of the furin cleavage site was detected when the virus was serially passaged 10 times in Vero cells. This contrasts with another recombinant live viral COVID-19 vaccine based on the VSV backbone^57^, where mutations arose at passage 9 in Vero E6 cells. One of these mutations occurred in the multibasic S1/S2 furin cleavage site and another generated a stop codon that resulted in a 24-amino acid truncation of the spike cytoplasmic tail. Truncation of the spike cytoplasmic tail was also reported when *wt* SARS-CoV-2^28^ or pseudotyped SARS-CoV-2^58, 59^ were propagated in tissue cultures. MV-014-212 virus shed from vaccinated AGMs had no mutations in the spike gene. Therefore, the chimeric spike gene in MV-014-212 appears to have a stable genotype in vitro and in vivo.

The vaccine profile of MV-014-212 remains unique among the current COVID-19 vaccines that have emergency use authorization or are in clinical development. MV-014-212 is administered intranasally, a needle-free route that offers potential advantages for global immunization. The intranasal route is similar to the natural route of infection of SARS-CoV-2 and generates both mucosal and humoral immune responses in AGMs without any adjuvant formulation. Modeling based on yields from production of phase 1 clinical study material projected a potential dose output of hundreds of millions of doses per annum in a modestly sized facility using high-intensity bioreactor systems. Mucosally delivered live attenuated vaccines such as MV-014-212 entail minimum downstream processing and have an anticipated low cost to produce. In addition, needle-free delivery reduces supply risks. Overall, MV-014-212 is well suited for domestic and global deployment as a primary vaccine or as a heterologous booster. MV-014-212 is currently being evaluated as a single-dose intranasal vaccine in a phase 1 clinical trial (NCT04798001).

## Materials and Methods

### Cells, viruses, and animals

Vero reference cell bank (RCB)1 (WHO Vero RCB 10-87) cells were grown in minimal essential medium (MEM, Gibco, Thermo-Fisher Scientific,) containing 10% fetal bovine serum (FBS, Corning) and 1x Corning Antibiotic/Antimycotic mix consisting of 100 IU/mL penicillin, 100 µg/mL streptomycin, 0.25 µg/mL amphotericin, with 0.085 g/L NaCI. RCB2 cells were derived from RCB1 and adapted to grow in serum-free media. RCB2 cells used in this study were grown in serum-free medium OptiPro (Gibco) supplemented with 4 mM of L-glutamine (Gibco). Both Vero cell lines were cultured at 37 °C, 5% CO_2_, with 95% humidity.

African green monkeys (*Chlorocebus aethiops*) were obtained from St. Kitts and were of indeterminate age, weighing 3-6 kg. The monkeys were screened and verified to be seronegative for RSV and SARS-CoV-2 by an RSV microneutralization assay and spike SARS-CoV-2 ELISA (BIOQUAL). Animals also underwent a physical examination by the veterinary staff to confirm appropriate health status prior to study. Each AGM was uniquely identified by a tattoo. One male and 3 females were assigned to the MV-014-212 and RSV groups. Two females and 1 male were assigned to the mock group. Cage-side observations included mortality, moribundity, general health, and signs of toxicity. Clinical observations included skin and fur characteristics, eye and mucous membranes, respiratory, circulatory, autonomic, and central nervous systems, somatomotor, and behavior patterns. The body weight of each monkey was recorded before the start of the dosing period and at each time of sedation. Consistent with the overall low levels of MV-014-212 replication in the respiratory tract of AGMs, no adverse events that were considered treatment-related were observed following inoculation with the vaccine. On Day 16 post vaccination one monkey inoculated with MV-014-212 died unexpectedly. Death occurred 4 days after the last NS and BAL surgical sample collection. A definitive determination of the cause of death could not be ascertained based on macroscopic or microscopic postmortem evaluations; however, there was no evidence that suggests the death was vaccine related. Moreover, the deceased animal had the lowest titer in NS samples compared to the other animals in this treatment group, with only one swab containing virus that was above the detection limit of the plaque assay (50 PFU/mL), and no detectable infectious virus in BAL at any of the time points evaluated.

Male and female K18-hACE2 Tg (strain #034860, B6.Cg-Tg[K18-ACE2]2Prlmn/J) mice were procured from The Jackson Laboratory (Bar Harbor, ME) and were approximately 8-10 weeks old at the time of vaccination.

The animal studies were conducted in compliance with all relevant local, state, and federal regulations and were approved by the BIOQUAL Institutional Animal Care and Use Committee (IACUC).

RSV Memphis37b was kindly provided by hVIVO (United Kingdom) and RSV TN12/11-19, by R.S. Peebles Jr (Vanderbilt University Medical Center).

### Plasmid construction

The recombinant MV-014-212 and derived viruses were cloned in the antigenome orientation in bacterial artificial chromosomes (BAC) under the control of the T7 polymerase promoter^24^. The BACs containing the recombinant MV-014-212 and MVK-014-212 sequences were constructed by restriction digestion and ligation from the DB1-QUAD and kRSV-DB1-QUAD plasmids (encoding the antigenome of an attenuated version of RSV with or without the *mKate* gene, respectively)^60^. The DNA sequence encoding the chimeric spike protein was designed to contain compatible cloning sites and it was synthesized by Twist Biosciences. The kRSV-DB1-QUAD plasmid and spike insert were digested with the enzymes AatII and SalI (NEB) and ligated with T4 DNA ligase (NEB) overnight at 16 °C. Stabl3 chemically competent cells (Invitrogen) were transformed with the ligation mix and selected for chloramphenicol resistance for 20-24 hours at 32 °C . MV-014-212 BAC was derived from the MVK-014-212 vector by removing the fragment between the KpnI and AatII restriction sites (∼7 kb containing the *mKate* gene) and replacing it with the corresponding fragment extracted from DB1-QUAD by restriction digestion and ligation. For all the constructs, the sequences of the entire encoded viruses were confirmed via Sanger sequencing. The construction of the plasmid rA2-mkate (aka, kRSV-A2) was described by Rostad et al^61^.

### Virus rescue and harvest

Vero cells were electroporated with the BAC encoding MV-014-212 (or the reporter virus) together with helper plasmids based on the pCDNA3.1 vector encoding the RSV proteins N, P, M2-1, and L, and the T7 polymerase under the control of a CMV promoter^24^. The cells were recovered in SFM-OptiPro medium supplemented with 4 mM glutamine and 10% fetal bovine serum (Hyclone) for 2 passages and then expanded in serum free medium with glutamine until CPE was extensive.

The recombinant viruses were harvested in Williams E medium (Hyclone) supplemented with sucrose phosphate glutamate buffer (SPG) or SPG alone by scraping the infected cells directly into the media. The lysate was vortexed for 15 seconds at maximum speed (3200 rpm) to release the viral particles and then flash frozen. One cycle of thawing and vortexing was performed to increase the release of virus before the stocks were aliquoted, flash-frozen, and stored at –70 °C until use.

### Plaque assay

Plaque assays for all the viruses used were performed in 24-well plates on Vero cells. Cells at 70% confluence were inoculated with 100 µL of 10-fold serial dilutions of viral samples (10^-1^ to 10^-6^). Inoculation was carried out at room temperature with gentle rocking for 1 hour before adding 0.75% methylcellulose (Sigma) dissolved in MEM supplemented with 10% FBS (Corning) and 1x Corning Antibiotic/Antimycotic mix. Cells were incubated for 4-5 days at 32 °C before fixing in methanol and immunostaining. For MV-014-212 and MVK-014-212, we used rabbit anti-SARS-CoV-2 spike polyclonal antibody (Sino Biological) and goat anti-rabbit HRP-conjugated secondary antibody (Jackson ImmunoResearch). For rA2-mKate2, the reagents used were goat anti-RSV primary antibody (Millipore) and donkey anti-goat HRP-conjugated secondary antibody (Jackson ImmunoResearch). In all cases, the viral plaques were stained with AEC (Sigma). The limit of detection is 1 PFU per well, corresponding to a minimum detectable titer of 100 PFU/mL.

### RNA sequencing

RNA from MV-014-212 samples was extracted using QIAamp^®^ Viral RNA Mini Kit following the protocol suggested by the manufacturer (Qiagen). The quality and concentration of the extracted RNA were evaluated by gel electrophoresis and UV spectrophotometry. The extracted RNA was used as the template for reverse transcription (RT) using Invitrogen SuperScript^®^ IV First-Strand Synthesis System using a specific primer or random hexamers. The cDNA 2nd strand was synthesized with the Platinum^TM^ SuperFi^TM^ PCR Master Mix. The purified PCR products were directly sequenced using the BigDye^®^ Terminator v3.1 Cycle Sequencing Kit (Applied Biosystems). The sequencing reactions were purified using Sephadex G-50 purification and analyzed on ABI 3730xl DNA Analyzer. The sequence traces were assembled using Sequencher software and the assembly was manually confirmed. The RNA sequencing for this study was performed by Avance Biosciences Inc., Houston TX.

### Western blot

Viruses and control recombinant SARS-CoV-2 spike protein (LakePharma) were denatured with Laemmli sample buffer (Alfa Aesar) by heating at 95 °C for 10 minutes. Proteins were separated by SDS-PAGE in a 4%-15% gradient gel and transferred to polyvinylidene fluoride (PVDF) membranes using a transfer apparatus according to the manufacturer’s protocol (BIO-RAD). After transfer, blots were washed in deionized water and probed using the iBind Flex system according to the manufacturer’s protocol (Invitrogen, Thermo-Fisher). Rabbit anti-SARS-CoV-2 spike (Sino Biological Inc, Beijing, China) was diluted in iBind solution (Invitrogen) at 1:1000. HRP-conjugated anti-rabbit IgG (Jackson ImmunoResearch) was diluted in iBind Solution at 1:5000. Blots were washed in deionized water and developed with ECL system (Azure Biosystems) according to manufacturer’s protocol. The blots were stripped with Restore Western Blot Stripping Buffer (Thermo-Fisher) and reprobed with goat anti-RSV polyclonal antisera (Sigma-Aldrich) and a monoclonal antibody specific for GAPDH (6C5) protein (Thermo-Fisher).

### Plaque assay for detecting virus shedding in AGMs

NS and BAL samples were collected and stored on ice until assayed for vaccine shedding by plaque assay. Vero cells were seeded in 0.5 mL per well at 1 x 10^5^ cells/mL in culture media in 24-well plates. The plates were incubated overnight at 37 °C in a humid incubator containing 5% CO_2_. The samples were diluted in Dulbecco’s modified Eagle medium (DMEM) without serum by adding 30 μL of nasal swab or BAL to 270 μL of DMEM. A total of six 10-fold serial dilutions were prepared in DMEM from 10^-1^ to 10^-6^. The media were removed from the 24-well plate and 100 μL of each dilution was added to duplicate wells of the 24-well plate of Vero cells. The plate was incubated at room temperature with constant rocking on a Rocker 35EZ, Model Rocker 35D (Labnet) for 1 hour. At the end of this incubation, 1 mL of methyl cellulose media (MEM supplemented with 10% fetal bovine serum, 1x antibiotic/antimycotic, and 0.75% methyl cellulose) was added to each well. The plate was incubated at 34 °C for 6 days in a humid incubator containing 5% CO_2_.

The plaques were visualized by immunostaining using RSV or SARS-CoV-2 antibodies. For immunostaining, the methyl cellulose media were aspirated, and the cell monolayers were washed with 1 mL of PBS at room temperature. The PBS was removed, and the cells were fixed by the addition of 1 mL of methanol to each well, and the plate was incubated at room temperature for 15 minutes. The methanol was removed, and cells washed with 1 mL of PBS followed by the addition of 1 mL Blotto solution (5% nonfat dried milk in Tris-buffered saline, Thermo-Fisher). The plates were incubated at room temperature for 1 hour. The Blotto solution was removed, and 0.25 mL of primary goat anti-RSV polyclonal antibodies (Millipore) diluted 1 to 500 in Blotto was added to RSV-infected cells. Cells infected with MV-014-212 were stained with primary rabbit anti-SARS-CoV-2 spike protein polyclonal antisera (Sino Biologicals, Beijing, China). The plates were incubated for 1 hour at room temperature with constant rocking. Primary antibodies were removed, and wells were washed with 1 mL Blotto solution.

For RSV-infected cells, 0.25 mL of donkey anti-goat HRP-conjugated polyclonal antisera (Jackson ImmunoResearch) diluted 1:250 in Blotto was added to each well. For MV-014-212– infected cells, goat anti-rabbit HRP-conjugated polyclonal antisera (Jackson ImmunoResearch) diluted 1:250 in Blotto was added to each well. The pate was incubated for 1 hour at room temperature with constant rocking. After incubation, the secondary antibodies were removed, and the wells washed with 1 mL of PBS. Developing solution was prepared by diluting AEC substrate 1:50 in 1x AEC buffer solution. A total of 0.25 mL of developing solution was added to each well and the plate was incubated at room temperature for 15 to 30 minutes with constant rocking until red immunostained plaques were visible. The developing reaction was terminated by rinsing the plate under tap water. The plaques were enumerated, and titers were calculated.

### RT-qPCR of SARS-CoV-2 subgenomic RNA for detecting shedding of challenge virus

The standard curve was prepared from frozen RNA stocks and diluted to contain 10^6^ to 10^7^ copies per 3 μL. Eight 10-fold serial dilutions of control RNA were prepared using RNAse-free water to produce RNA concentrations ranging from 1 to 10^7^ copies/reaction.

The plate was placed in an Applied Biosystems 7500 Sequence detector and amplified using the following program: 48 °C for 30 minutes, 95 °C for 10 minutes followed by 40 cycles of 95 °C for 15 seconds, and 1 minute at 55 °C. The number of copies of RNA per mL of sample was calculated based upon the standard curve.

Total RNA from tissues was extracted using RNA-STAT 60 (Tel-test “B”)/ chloroform followed by precipitation of the RNA and resuspension in RNAse-free water. To detect SARS-CoV-2 sgRNA, a primer set and probe were designed to detect a region of the leader sequence and *E* gene RNA from SARS-CoV-2. The *E* gene mRNA is processed during replication to contain a 5’ leader sequence that is unique to sgRNA (not packaged into the virion) and therefore can be used to quantify sgRNA. A standard curve was prepared using known quantities of plasmid DNA containing the *E* gene sequence, including the unique leader sequence, to produce a concentration range from 1 to 10^6^ copies/reaction. The PCR reactions were assembled using 45 μL master mix (Bioline) containing 2x buffer, Taq-polymerase, reverse transcriptase, and RNAse inhibitor. The primer pair was added at 2 μM, and 5 μL of the sample RNA was added to each reaction in a 96-well plate. The PCR reactions were amplified in an Applied Biosystems 7500 Sequence detector using the following conditions: 48 °C for 30 minutes, 95 °C for 10 minutes followed by 40 cycles of 95 °C for 15 seconds, and 1 minute at 55 °C.

Primers / Probe sequences are shown below:

SG-F: CGATCTTGTAGATCTGTTCCTCAAACGAAC
SG-R: ATATTGCAGCAGTACGCACACACA
FAM-ACACTAGCCATCCTTACTGCGCTTCG-BHQ

### TCID50 assay to detect shedding of SARS-CoV-2 in BAL and NS of AGMs after challenge

Vero TMPRSS2 cells (obtained from Adrian Creanga, Vaccine Research Center-NIAID) were plated at 25,000 cells/well in DMEM + 10% FBS + gentamicin and the cultures were incubated at 37 °C, 5.0% CO_2_. Cells were 80%-100% confluent the following day. Medium was aspirated and replaced with 180 μL of DMEM + 2% FBS + gentamicin. Twenty (20) μL of sample was added to the top row in quadruplicate and mixed using a P200 pipettor 5 times. Using the pipettor, 20 μL was transferred to the next row, and repeated down the plate (columns A-H), representing 10-fold dilutions. Positive (virus stock of known infectious titer in the assay) and negative (medium only) control wells were included in each assay set-up. The plates were incubated at 37 °C, 5.0% CO_2_ for 4 days. The cell monolayers were visually inspected for CPE. The TCID_50_ value was calculated using the Read-Muench formula.

### SARS-CoV-2 total IgG ELISA for AGM sera

MaxiSorp immuno plates (Thermo-Fisher) were incubated overnight at 4 °C with 100 μL of 0.65 μg/mL of SARS-CoV-2 spike prepared in PBS (Pre-S SARS-CoV-2 spike, Nexelis). The protein solution was removed, and the plate was washed 4 times with 250 μL of PBS supplemented with 0.05% Tween 20 (PBST). Blocking solution (PBST containing 5% nonfat dried milk) was added at 200 μL per well and the plate was incubated for 1 hour at room temperature. A SARS-CoV-2 spike-specific IgG (Nexelis) was diluted in blocking solution and used as a standard. Negative control serum was diluted 1:25 in blocking solution. Serum samples were diluted at 1:25 followed by eight 2-fold serial dilutions in blocking solution. The blocking solution was removed from the plate and the wells washed once with 250 μL of PBST followed by addition of 100 μL of the diluted serum samples and controls, and the plate was incubated for 1 hour at room temperature. The plate was washed 4 times with 250 μL PBST and 100 μL of HRP-conjugated goat anti-monkey IgG antibody (PA1-8463, Thermo-Fisher) diluted in blocking solution was added to each well following the last wash step. The plate was incubated for 1 hour at room temperature and then washed 4 times in 250 μL PBST. Developing solution containing 3, 3’, 5, 5’ - Tetramethylbenzidine (TMB) substrate (1-Step Ultra TMB-ELISA Substrate Solution, Thermo-Fisher) was added to each well and the plate was incubated at room temperature for 30 minutes to allow the color to develop. The colorimetric reaction was terminated by the addition of 100 μL of ELISA Stop Solution (Invitrogen). The absorbance at 450 nm and 650 nm was read by spectrophotometry using a SpectraMax iD3 microplate reader (Molecular Devices).

### SARS-CoV-2 IgA ELISA for AGM nasal swabs

Purified pre-fusion SARS-CoV-2 spike antigen (LakePharma) was adsorbed onto 96-well MaxiSorp immuno microplate (Thermo-Fisher). The positive control was a serum pool from 3 COVID-19 convalescent individuals (Nexelis). Total IgA purified from human serum was used as a standard (Sigma-Aldrich). To generate the IgA standard curve anti-human IgA capture antibodies, Mab MT57 (Mabtech), were absorbed on plates instead of spike antigen. Following incubation, the microplate was washed 4 times with 250 µL PBST and blocked with 1% BSA in PBST. Purified human IgA standard, controls, or sample dilutions were then added and incubated in the coated microplate to allow binding. The plates were washed and a biotinylated goat anti-human IgA antibody (Mabtech) with cross-reactivity to monkey antibodies was added to all wells. Excess biotinylated anti-IgA antibody was removed by washing and streptavidin-conjugated HRP (Southern Biotech) was added. TMB was added and color development was stopped by addition of stop solution from Invitrogen. The absorbance of each well was measured at 450 nm. The standard total IgA antibody assayed on each test plate was used to calculate the concentration of IgA antibodies against spike protein in the AGM samples expressed in the arbitrary units ELU/mL. The measurements were performed in duplicate and average values are reported with standard deviations.

### Microneutralization assay

Heat-inactivated sera from the AGMs were diluted serially in MEM with nonessential amino acids (Gibco) and antibiotics/antimycotic. All experiments were done in duplicate. 200 PFU of the desired reporter virus were added to each dilution and incubated at room temperature for 1 hour. Confluent RCB1 cells grown in a clear-bottom black 96-well plate (Grenier) were infected with the serum-virus mixes and centrifuged (spinoculated) at 1800 x g for 30 minutes at 20 °C. The plates were incubated for 20 hours at 37 °C and 5% CO_2_. The fluorescent foci in each well were counted using a Celigo Image Cytometer (Nexcelom) and converted to % inhibition using the formula below:

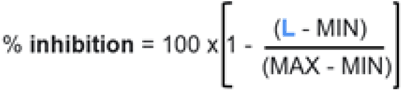

where MIN is the average number of foci obtained in the control wells with only cells (no virus) and MAX is the average number of foci from the wells in the control wells with only virus (no serum). L is the number of foci in the sample wells. The resulting curves of inhibition vs. dilution of the sera were fitted using non-linear regression, option “[inhibitor] vs normalized response-variable slope” in GraphPad Prism (version 9.0.0). From the fitting, half-maximal inhibitory concentration (IC_50_) was obtained and NT_50_ was calculated as the reciprocal of IC_50_.

### Pseudovirus neutralization assay

SARS-CoV-2 spike ΔCT protein-bearing vesicular stomatitis virus (VSV) pseudotyped particles in which the VSV glycoprotein *G* gene was replaced by the luciferase reporter gene were purchased from Nexelis, and the assay was performed at Meissa. Starting with a dilution of 1/25, serial 2-fold dilutions of heat-inactivated AGM sera were incubated with the pseudotyped viral particles (3 x 10^5^ RLU/well) at 37 °C for 1 hour, after which the mixes were used to infect Vero E6 monolayers in white 96-well plates. The cells were incubated with the infection mixes for 20 ± 2 hours at 37 °C with 5% CO_2_. The medium was then discarded, and the cells were lysed in ONE-GloTM EX Luciferase Reagent (Promega). The lysis and luciferase reactions were allowed to proceed for 3 minutes at room temperature with shaking at 600 rpm and luminescence was read with a SpectraMax i3D plate reader. The values of luminescence were converted to percentage of inhibition using the equation shown in the previous paragraph where MIN is the average of reads obtained in the cell-only wells and MAX is the average of the reads from the wells of the pseudovirus-only control. L is the sample luminescence value. The inhibition vs dilution curves were fitted using nonlinear regression, option “[inhibitor] vs normalized response-variable slope” in GraphPad Prism (version 9.0.0). From the fitting IC_50_ was obtained and NT_50_ was calculated as the reciprocal of IC_50_.

### Plaque reduction neutralization test

A conventional 50% plaque-reduction neutralization test (PRNT_50_) was performed to quantify the serum-mediated virus suppression as previously reported^39^. Briefly, individual sera were 2-fold serially diluted in culture medium with a starting dilution of 1:40 (dilution range of 1:40 to 1:1280). The diluted sera were incubated with 100 PFU of USA-WA1/2020 or mutant SARS-CoV-2. After 1 hour of incubation at 37 °C, the serum-virus mixtures were inoculated onto a monolayer of Vero E6 cells pre-seeded on 6-well plates on the previous day. A minimal serum dilution that suppresses >50% of viral plaques is defined as PRNT_50_.

All recombinant SARS-CoV-2s with spike mutations^40^ were prepared on the genetic background of an infectious cDNA clone derived from clinical strain USA-WA1/2020^62^.

### Spike-specific IgG1 and IgG2a ELISA

Serum samples from mice were collected on Day –21 and on Day 28 post vaccination to quantify the levels of SARS-CoV-2 spike-specific IgG1 and IgG2a antibodies by ELISA. Purified perfusion-stabilized SARS-CoV-2 spike protein (SARS-CoV-2/human/USA/WA1/2020, from LakePharma) was diluted to 1 µg/mL in PBS and 100 μL was added to each well of a MaxiSorp immuno plate (Thermo-Fisher) and incubated overnight at 4 °C. The plate was washed 4 times in PBST (PBS + 0.05% Tween 20) and 100 μL of blocking solution (PBST + 2% BSA) was added to each well, and the plate was incubated for 1 hour at room temperature. Serum dilutions were prepared in blocking solution with the first dilution at 1:25 for the IgG1 assay or 1:10-1:100 for the IgG2a assay. SARS-CoV-2 spike IgG1 (Sino Biological) or anti-spike-RBD- mIgG2a (InvivoGen) were diluted in blocking solution and used as standards for the assay. The blocking solution was removed and 100 μL of diluted antibody added to each well. The plate was incubated at room temperature for 1 hour and then washed 4 times in PBST using the plate washer. Then, 100 μL of HRP-conjugated goat-anti-mouse IgG1 (Thermo-Fisher) or HRP-conjugated goat-anti-mouse IgG2a (Thermo-Fisher) secondary antibodies diluted 1:32,000 and 1:1000, respectively, were added to each well and the plate was incubated at room temperature for 1 hour. The plate was washed 4 times in PBST. 100 μL of 1-step ultra TMB-ELISA substrate solution (Thermo-Fisher) was added to each well and the plate incubated for 30 minutes with constant rocking on an orbital shaker. After the incubation period, 100 μL of stop solution (Invitrogen) was added to each well and the plate read on a Spectramax id3 plate reader (Molecular Devices) at 450 nm and 620 nm.

### ELISPOT of splenocytes from MV-014-212–vaccinated hACE2-mice

Spleens from vaccinated ACE2 mice were collected on Day 28 post inoculation and stored in DMEM containing 10% FBS on ice until processed. The spleens were homogenized on a sterile petri dish containing medium. The homogenate was filtered through a 100-μm cell strainer and the cell suspension transferred to a sterile tube on ice. The cells were collected by centrifugation at 200 x g for 8 minutes at 4 °C. The supernatant was removed and residual liquid on the edge of the tube blotted with a clean paper towel. Red blood cells were lysed by resuspending the cell pellet in 2 mL of ACK lysis buffer (155 mM ammonium chloride, 10 mM potassium bicarbonate, 0.1 mM ethylenediaminetetraacetic acid [EDTA]) and incubating the samples at room temperature for approximately 5 minutes. PBS was added at 2x to 3x the volume of cell suspension and cells were collected by centrifugation at 200 x g for 8 minutes at 4 °C. The cell pellet was washed twice in PBS and the cells collected by centrifugation at 200 x g for 8 minutes at 4 °C. The supernatant was removed, and the pellet resuspended in 2 mM L-Glutamine CTL-Test Media (Cell Technology Limited). The suspension was filtered through a 100-μm cell strainer into a new 15-mL conical tube and the cells counted using a hemocytometer and resuspended at the appropriate cell concentration. Cells were maintained at 37 °C in a humidified incubator with 5% CO_2_ until used in the ELISpot assay.

The ELISpot assay was performed using a mouse IFNγ/IL-5 Double-Color ELISPOT assay kit (Cell Technology Limited). Murine IFNγ/IL-5 capture solution and 70% ethanol was prepared according to the manufacturer’s protocol. The membrane on the plate was activated by addition of 15 μL of 70% ethanol to each well. The plate was incubated for less than 1 minute at room temperature followed by addition of 150 μL PBS. The underdrain was removed to drain the solution in the wells and each well was washed twice with PBS. Murine IFNγ/IL-5 capture solution (80 μL) was added to each well and the plate was sealed with parafilm and incubated at 4 °C overnight. The capture solution was removed, and the plate washed once with 150 μL PBS. A peptide pool containing peptides of 15 amino acids in length that span the SARS-CoV-2 spike protein (PepMix™ SARS-CoV-2 spike Glycoprotein, JPT Peptide Technologies) were prepared at 10 mg/mL and 100 μL was added to each well. A positive control containing Concanavalin A (Con A) mitogen (10 μg/mL) was added to a separate reaction mixture. The splenocytes were mixed with CTL-Test™ Medium (Cell Technology Limited) to yield a final cell density of 3,000,000 cells/mL and 100 μL/well were added to the plate using large orifice tips. The plate was incubated at 37 °C in a humidified incubator containing 9% CO_2_ for 24 hours. The plates were washed twice with PBS and then twice with 0.05% Tween-PBS at a volume of 200 μL/well for each wash followed by addition of 80 μL/well anti-murine IFNγ/IL-5 detection solution. The plates were incubated at room temperature for 2 hours. The plate was washed 3 times with PBST at 200 μL/well for each wash followed by the addition of 80 μL/well of tertiary solution. The plates were incubated at room temperature for 1 hour. The plate was washed twice with PBST, and then twice with 200 μL/well of distilled water. Blue Developer Solution was added at 80 μL/well and the plate was incubated at room temperature for 15 minutes. The plate was rinsed 3 times in tap water to stop the developing reaction. After the final wash, red developer solution was added at 80 μL/well and the plate was incubated at room temperature for 5-10 minutes. The plate was rinsed 3 times to stop the developing reaction. The plate was air-dried for 24 hours face-down on paper towels on the bench top. The spots on the plate representing splenocytes expressing IFNγ (red) or IL-5 (blue) were quantified using the CTL-immunospot plate reader (ImmunoSpot 7.0.23.2 Analyzer Professional DC\ImmunoSpot 7, Cellular Technology Limited) and software (CTL Switchboard 2.7.2).

### Data availability statement

Source data are provided with this paper. The sequences of the antigenomes of MV-014-212 and MVK-014-212 are deposited in GenBank with accession numbers MZ695841 and MZ695842.

## Acknowledgement

We thank Anthony Cook, Renita Brown, Tammy Putmon-Taylor, Tracey-Ann Campbell, Shanai Browne, Zack Flinchbaugh, Elyse Teow and Jason Velasco at Bioqual for animal care support and technical expertise with our African green monkey study. We thank Douglas Haney for biostatistics support and Nexelis for sharing pseudovirus neutralization assay and reagents. We thank Tina V. Hartert from the Peebles’ lab for the TN 12/11-19 RSV strain and hVIVO for the Memphis 37b RSV strain. We are grateful to Janelle Muranaka for technical and logistical support.

## Author Contributions

M.F.T., R.J., C.C.S., M.L.M., and R.S.T. participated in study concept/design. M.F.T., R.J., A.S.P., A.G., D.W., S.P., X.C., J.G., T.O., D.V., S.K., L.P., H.A., M.H.B., R.S.P., Y.L., X.X., and R.S.T. assisted with data acquisition. M.F.T. and R.J. provided statistical analyses. M.F.T., R.J. P-Y.S., M.L.M., and R.S.T interpreted the data. The manuscript was written by M.F.T., R.J., and R.S.T. with contributions from all authors. All authors provided final approval for submission.

