## Supplemental Material for "One mucosal administration of a live attenuated recombinant COVID-19 vaccine protects nonhuman primates from SARS-CoV-2"

### Supplementary Information

**Fig. S1: Other vaccine candidates designed in this study.** The sequences of the C-termini of the candidates show the different positions of the junction between the RSV F protein cytoplasmic tail (red font) and the SARS-CoV-2 spike protein transmembrane domain (blue font). For reference, the full-length sequences of the C-termini of spike and F are shown above and below the candidates, respectively. The candidates were designed to contain an *mKate2* gene as a fluorescent marker to follow rescue and propagation in culture. The 7 candidates listed in this figure (and table below) were evaluated by their ability to be rescued (defined as the generation of red fluorescent foci) and to grow to titers of $10^5$ PFU/mL or higher. MV-014-212 was chosen to pursue further investigation. RSV, respiratory syncytial virus.

```
                       TM
IDLQELGKYEQYIKWPWYIWLGFIAGLIAIVMVTIMLCCMTSCCSCLKGCCSCGSCCKFDEDDSEPVLKGVKLHYT                 SARS-CoV-2 spike

IDLQELGKYEQYIKWPWYIWLGFIAGLIAIVMVTIMLCCMTSCCSCLKGCCSCGSCC-----------------KARSTPVTLSKDQLSGINNIAFSN    MV-014-210
IDLQELGKYEQYIKWPWYIWLGFIAGLIAIVMVTIMLCCMTSCCSCLKGCCSCGSCCKFDEDD---------KARSTPVTLSKDQLSGINNIAFSN    MV-014-211
IDLQELGKYEQYIKWPWYIWLGFIAGLIAIVMVTIMLCCMTSCCSCLKGCCSCGSCCKFDEDDGLLLYCKARSTPVTLSKDQLSGINNIAFSN    MV-014-212
IDLQELGKYEQYIKWPWYIWLGFIAGLIAIVMVTIMLCCMTSCCSCLK------------------------GLLLYCKARSTPVTLSKDQLSGINNIAFSN    MV-014-220
IDLQELGKYEQYIKWPWYIWLGFIAGLIAIVMVTIMLCCMTSCCSCL-----------------------------------KARSTPVTLSKDQLSGINNIAFSN    MV-014-230
IDLQELGKYEQYIKWPWYIWLGFIAGLIAIVMVTIMLCCMTSCCSCLK------------------------------YCKARSTPVTLSKDQLSGINNIAFSN    MV-014-240
IDLQELGKYEQYIKWPWYIWLGFIAGLIAIVMVTIMLCCMTSCCSCLKKGCCSCGSCCKFDEDDSEPVLKGVKLHYT                MV-014-300

             SQVNEKINQSLAFIRKSDELLHNVNAGKSTTNIMITTIIIVIIVILLSLIAVGLLLYCKARSTPVTLSKDQLSGINNIAFSN    RSV F
                                                     TM
```

| Vaccine candidate | Rescue | Achieved titers ≥ $10^5$ PFU/mL |
|---|---|---|
| **MV-014-210** | **Y** | **Y** |
| **MV-014-211** | **Y** | **N.D.** |
| **MV-014-212** | **Y** | **Y** |

| | | |
|---|---|---|
| **MV-014-220** | **Y** | **N.D.** |
| **MV-014-230** | **N** | **N** |
| **MV-014-240** | **N** | **N** |
| **MV-014-300** | **Y** | **N** |

Y: yes

N: no

N.D.: Not determined

**Fig. S2: Schematic of spike and its glycosylation sites.** Based on its amino acid sequence, the calculated molecular weight of the chimeric spike protein monomer in MV-014-212 prior to glycosylation is approximately 143 kDa, (https://www.bioinformatics.org/sms/prot_mw.html). Wild type spike has a calculated molecular weight of 141.2 kDa without counting the glycosylation, whereas the apparent weight is around 180 kDa[1]. We expect the chimeric protein encoded by MV-014-212 to be similarly glycosylated and show a similar apparent size in an SDS-PAGE gel.

Upon proteolysis at the S1-S2 junction, MV-014-212 spike would give rise to two subunits of 683 and 607 amino acids, respectively. Both these subunits would be glycosylated[2] and likely be indistinguishable in SDS-PAGE as observed by Peacock et al[3] (2021). The figure shows a schematic of spike and its glycosylation sites (short black bars) as reported by Watanabe et al[2]. S1, subunit 1; S2, subunit 2; SDS-PAGE, sodium dodecyl-sulfate polyacrylamide gel electrophoresis.

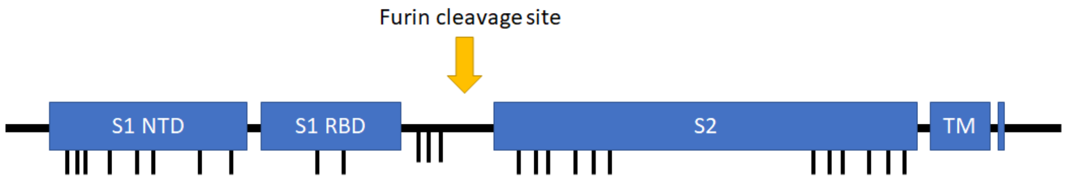

**Fig. S3: Design of the genetic stability experiment.** A stock of MV-014-212 (passage 0) was used to infected three independent T75 flasks of subconfluent Vero cells (P1), establishing three lineages. The P1 flasks were harvested and used for sequential passage for 10 total passages. RNA was extracted from virus in Passages 0 and 10 (3 lineages), and the coding regions of the genome were sequenced by Sanger sequencing,

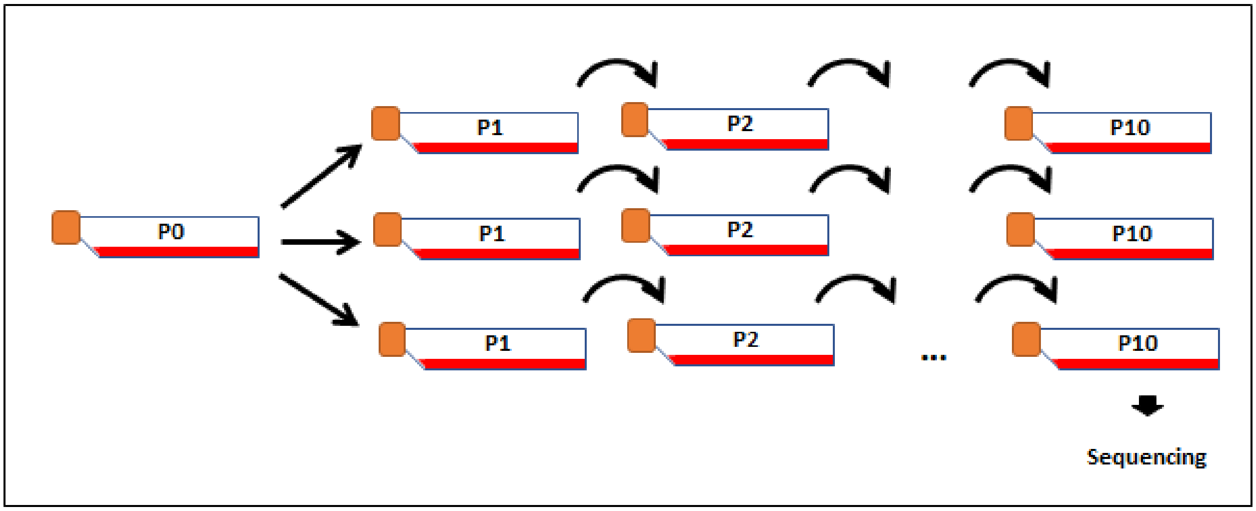

**Fig. S4: Shedding kinetics in individual AGMs after vaccination.** The panels on the left correspond to the AGMs vaccinated with MV-014-212 and the panels on the right are AGMs in the control group inoculated with RSV. AGM, African green monkey; BAL, bronchoalveolar lavage, RSV, respiratory syncytial virus.

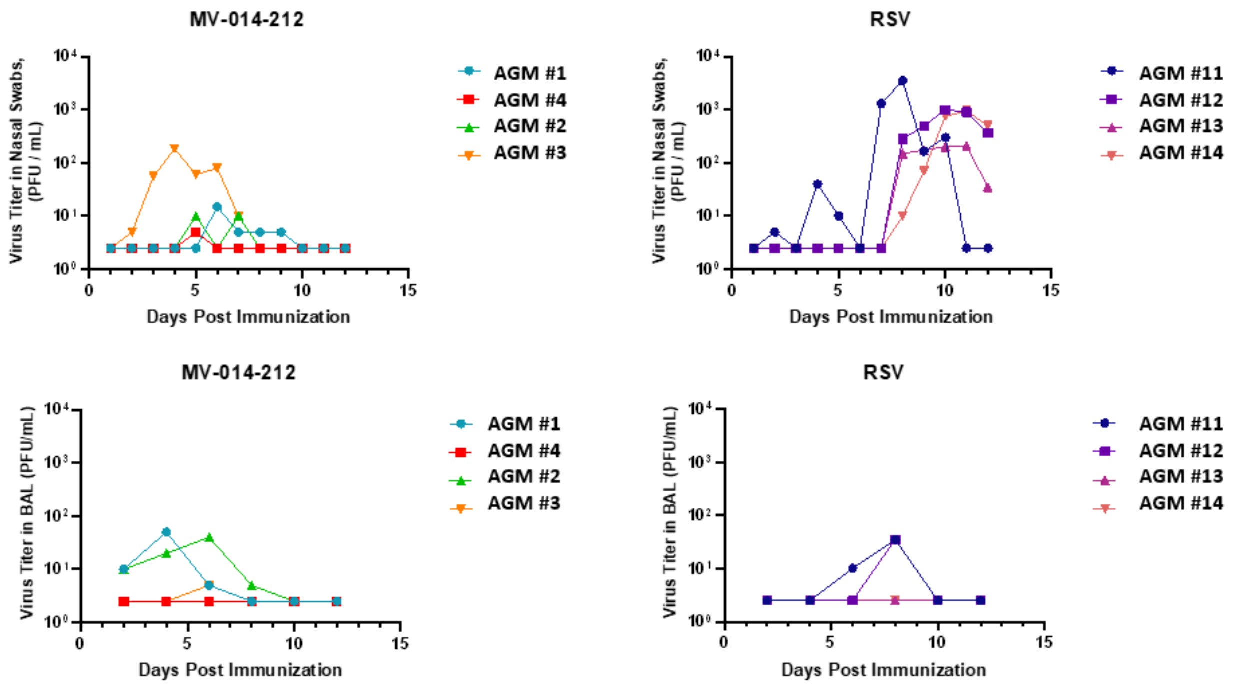

**Fig. S5:** Two independent experiments were performed in cotton rats. In experiment 1, cotton rats (*n* = 5 per group) were inoculated with 1 x $10^5$ PFU of biologically derived RSV TN 12/11-19 (TN12)[4], Memphis 37b (M37)[5], or recombinant A2 (rA2) RSV strains. On Days 3, 5, and 7 cotton rat nasal and lung tissues were homogenized in HBSS + 10% SPG for titer determination by plaque assay. In this experiment only Day 5 nasal and lung tissues were collected for the rA2 group. In experiment 2, cotton rats (*n* = 6) were inoculated with 5 x $10^5$ PFU of rA2. On Days 2, 5, and 7 cotton rat nasal and lung tissues were homogenized in HBSS +10% SPG for titer determination by plaque assay. Plaque assay was performed in HEp-2 cells using clarified nasal and lung homogenates diluted in EMEM. Plaques were visualized by immunostaining with RSV polyclonal antibodies in experiment 1 and by crystal violet staining in experiment 2. Panel **a** shows replication kinetics of TN12, M37, and rA2 in the nose. Panel **b** compares nasal titers of TN12, M37, and rA2 on Day 5 of experiment 1. Panel **c** shows replication kinetics of biological TN12 and Memphis 37 and rA2 in the lungs. Panel **d** compares lung titers on Day 5 of experiment 1 for TN12, M37, and rA2. The result showed that the rA2 nose titers were comparable to biologically derived TN12 and M37 but rA2 lung titers were approximately 2 logs lower compared with biologically derived RSV strains. EMEM, Eagle's minimum essential medium; Hank's balanced salt solution; RSV, respiratory syncytial virus; SPG, sucrose phosphate glutamate.

**a.**

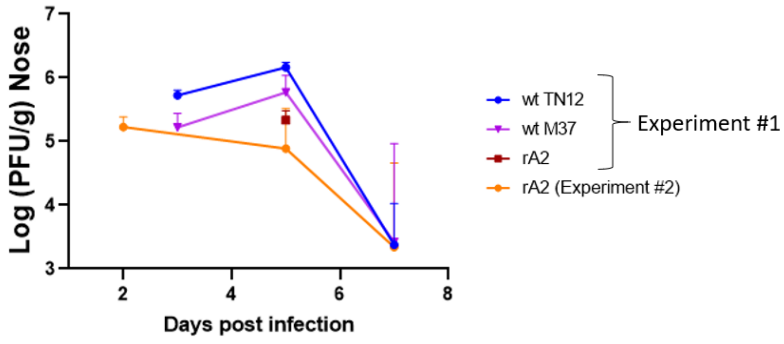


**b.**

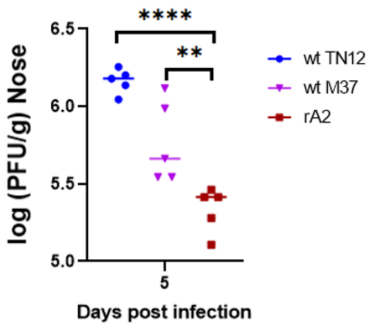

**c.**

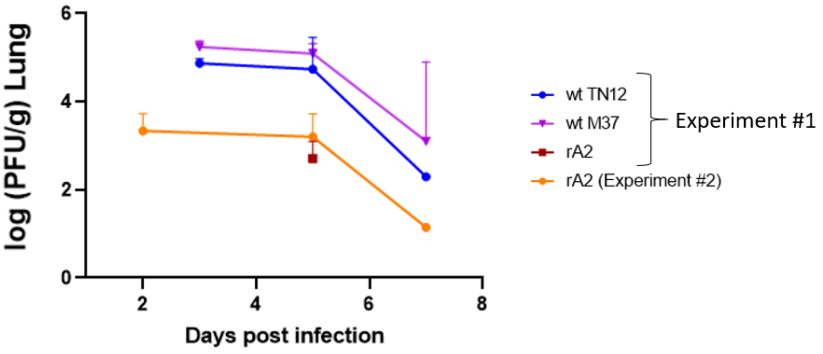
log (PFU/g) Lung
Days post infection
0
2
4
6
8
wt TN12
wt M37
rA2
rA2 (Experiment #2)
Experiment #1

**d.**

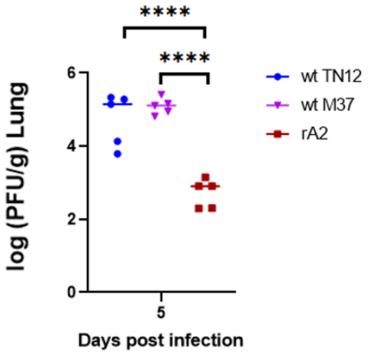
****
****
log (PFU/g) Lung
Days post infection
0
2
4
6
5
wt TN12
wt M37
rA2

**Fig. S6: SARS-CoV-2 shedding kinetics in individual AGMs after challenge.** The panels on the left correspond to the AGMs vaccinated with MV-014-212, the panels in the center are AGMs in the control group inoculated with RSV, and the panels on the right are the AGMs in the control group that was mock-inoculated. AGM, African green monkey; RSV, respiratory syncytial virus.

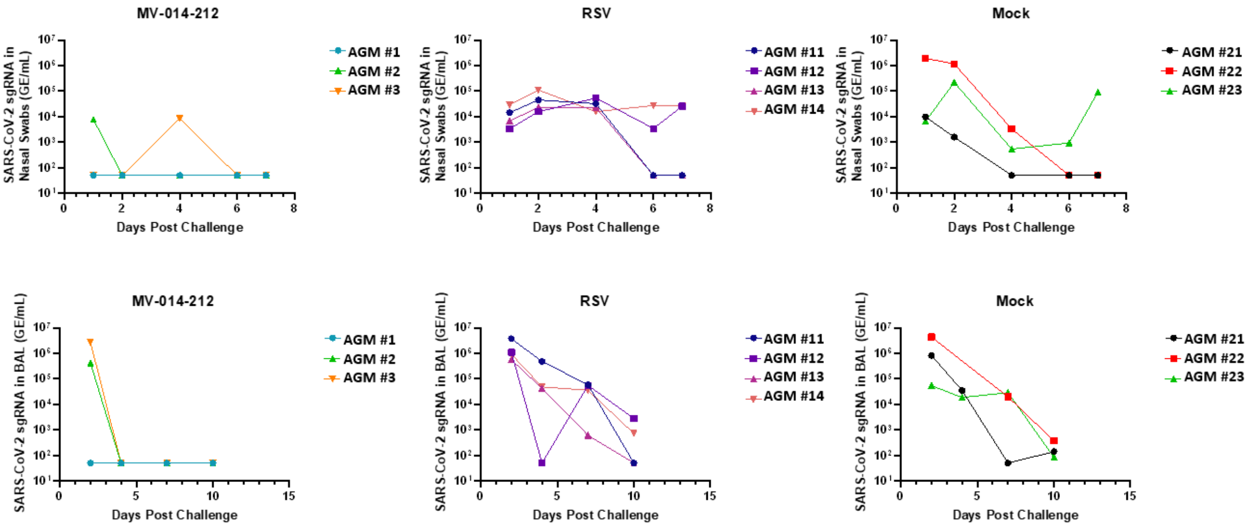

**Fig. S7: TCID₅₀ of SARS-CoV-2 shedding.** The panel on the top left corresponds to samples from NS and the panel on the top right is from BAL samples. Lines indicate mean and the error bars are SD. The bottom panels are time courses for the individual animals in the same experiment. BAL, bronchoalveolar lavage; NS, nasal swabs; RSV, respiratory syncytial virus; SD, standard deviation; $TCID_{50}$, median tissue culture infectious dose.

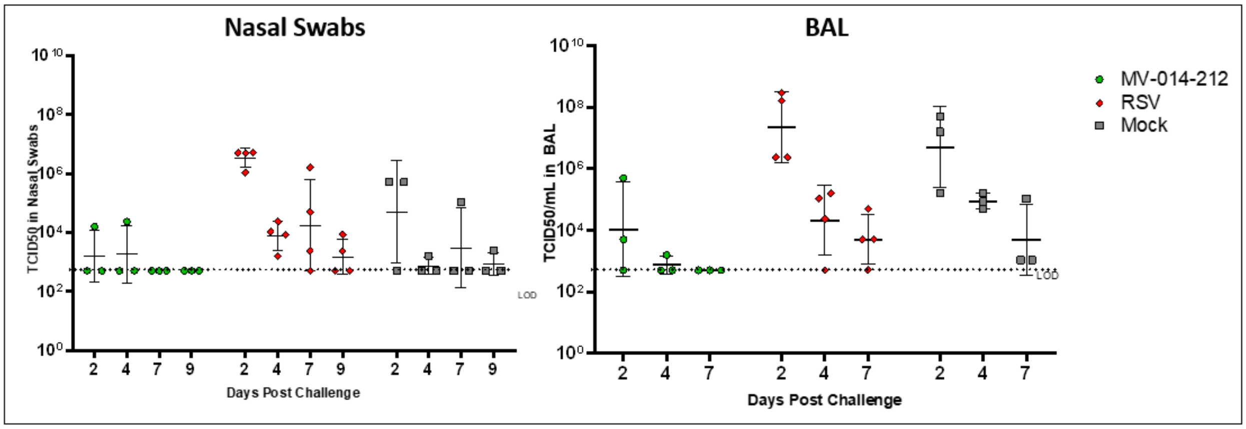


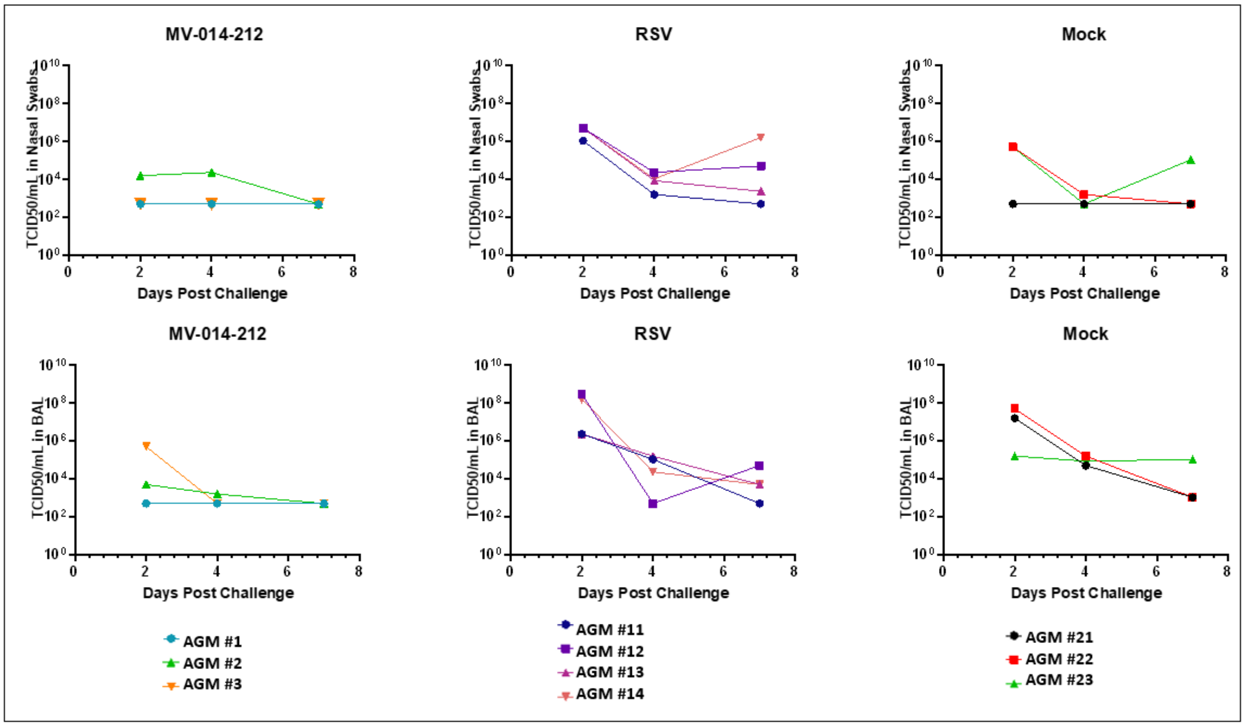

**Fig. S8: Pseudovirus neutralization assay.** Neutralizing titers ($NT_{50}$) in serum of AGMs immunized either with MV-014-212 or the negative controls, RSV or mock, obtained from a luciferase-based pseudovirus neutralization assay using VSV-spike pseudoparticles (Nexelis). "Conv." is a pool of 3 human convalescent sera provided by Nexelis with an informed titer of 1859 IU/mL. $NT_{50}$ was obtained from fitting inhibition curves as described in Methods. RSV, respiratory syncytial virus.

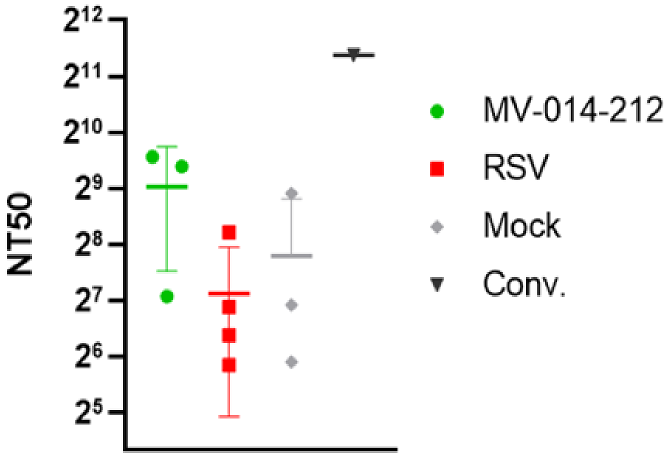

**Table S1: Shedding and immunologic data of individual AGMs vaccinated with MV-014-212**

| | AGM #1 | AGM #2 | AGM #3 | AGM #4 |
|---|---|---|---|---|
| Peak shedding of vaccine (MV-014-212) in NS (PFU/mL) | 15 | 10 | 185 | 5 |
| Peak shedding of vaccine (MV-014-212) in BAL (PFU/mL) | 50 | 40 | 5 | 2.5 (LOD) |
| Peak shedding of SARS-CoV-2 (challenge) in NS (GE/mL) | 50 (LOD) | $8.5\ 10^3$ | $7.7\ 10^3$ | – |
| Peak shedding of SARS-CoV-2 (challenge) in BAL (GE/mL) | 50 (LOD) | $2.8\ 10^6$ | $4.2\ 10^5$ | – |
| Peak shedding of SARS-CoV-2 (challenge) in NS ($\log(TCID_{50})$/mL) | <2.7 | 4.4 | <2.7 | – |
| Peak shedding of SARS-CoV-2 (challenge) in BAL ($\log(TCID_{50})$/mL) | <2.7 | 3.7 | 5.7 | – |
| Serum $NT_{50}$ (microneutralization assay) | 139.4 | 114.8 | 42.5 | – |
| Serum $NT_{50}$ (pseudo virus assay) | 757.6 | 670.7 | 134.9 | – |
| Fold change spike-specific IgA in NS | 6.7 | 14.0 | 7.2 | – |
| Spike-specific IgG in serum (ELU/mL) | 5010.2 | 2174.0 | 399.0 | – |

AGM, African green monkey; BAL, bronchoalveolar lavage; Ig, immunoglobulin; LOD, limit of detection; NS, nasal swab; $NT_{50}$, neutralizing titers; $TCID_{50}$, median tissue culture infectious dose.
